# Bacterial Strain Identity and Community Composition Drive the Seed-to-Seedling Microbiota Assembly

**DOI:** 10.64898/2026.02.02.703190

**Authors:** Gontran Arnault, Coralie Marais, Anne Préveaux, Martial Briand, Samuel Jacquiod, Alain Sarniguet, Matthieu Barret, Marie Simonin

## Abstract

The seed-to-seedling transition is a key step of plant microbiota assembly, where the seed microbiota meets the soil microbiota. Seed germination triggers profound physiological changes, including exudate release and oxidative bursts, which generate selective pressures shaping microbial survival and interactions. Here, we investigated how host selection and community composition influence bacterial colonization during this transition using a Synthetic Community (SynCom) approach on common bean (*Phaseolus vulgaris*) seeds. Specifically, we tested the importance of microbial interactions by applying 30 bacterial SynComs spawning a phylogenetic diversity gradient.

Seedling colonization success was primarily determined by strain identity but was strongly modulated by the SynCom context. Strains that were highly effective colonizers when inoculated alone often exhibited reduced transmission within SynComs, indicating that microbial interactions can either inhibit or facilitate colonization. Random forest models confirmed that colonization outcomes could not be predicted from single-strain performance alone. Instead, both phylogenetic relatedness and metabolic similarity among SynCom members emerged as key predictors of strain success, supporting the competition–relatedness hypothesis and highlighting the importance of community-dependent effects in shaping colonization success. Genomic analyses identified microbial traits linked to efficient seedling colonization, notably amino acid transport and metabolism, stress tolerance, ROS detoxification, and biofilm formation, highlighting the strong selective pressures acting during the seed-to-seedling transition. Besides key adaptative traits, our findings underscore the importance of microbial interactions to survive and colonize seedlings, bringing a novel perspective for the successful engineering of seed and seedling microbiota.

## Introduction

Plants are associated with many microorganisms that influence their health and productivity (Arnault et al., 2023; Vandenkoornhuyse et al., 2015; Vannier et al., 2019). Engineering plant-associated microbiota has therefore emerged as a promising strategy to improve plant performance and resilience (Afridi et al., 2022). Indeed, synthetic communities (SynComs) inoculations (Vorholt et al., 2017; Mehlferber et al., 2024) can modulate plant growth (Kaur et al., 2022), limit disease symptoms (Yin et al., 2022) and enhance drought stress tolerance (Armanhi et al., 2021) in controlled growth conditions. However, translating these benefits to the field remains challenging (Russ et al., 2023). This transposability problem often reflects a lack of holistic understanding of microbiota assembly processes (Joubert et al., 2025). Indeed, most studies focus on the impact of SynComs on plant phenotype, without assessing the colonization success of the inoculated microorganisms. A better characterization of SynCom dynamics after inoculation is therefore needed, particularly regarding their ability to colonize plant tissues and the complex microbial interactions that shape community assembly (Russ et al., 2023; Hassani et al., 2018). The limited deployment of SynComs in field conditions can be attributed to several key constraints: (i) the large quantity of microbial cells required for effective plant colonization, (ii) our still limited understanding of the microbial processes governing successful colonization of plant compartments and (iii) the often low competitiveness of lab-assembled SynComs against well-adapted native microbial communities under local environment conditions (Joubert et al., 2025; Russ et al., 2023). These challenges highlight the need for alternative delivery strategies and more targeted microbial interventions capable of overcoming ecological and practical barriers.

To address these constraints, plant compartments offering both ecological leverage and practical feasibility are increasingly being explored. Among them, the seed microbiota and its engineering is being recognized as a particularly promising approach (Joubert et al., 2025). Indeed, seed microbiota engineering is a potential way to deliver plant-beneficial microorganisms using a limited amount of inoculum (Rocha et al., 2019; Arnault et al., 2024). Also, seeds represent the first microbial habitat of the plant life cycle and therefore may influence the assembly of the microbial communities in the subsequent plant compartments (Ridout et al., 2019; Debray et al., 2021). The transition from seed-to-seedling involves profound morphological and physiological changes. This phase marks the coalescence (*i.e* the mixing between two distinct communities) between the seed and the soil microbiota, particularly in the spermosphere, where seeds release a plethora of exudates and undergo oxidative bursts (Scarafoni et al., 2013; Nelson 2018; Joubert et al., 2025; Bailly 2023). These exudates create a nutrient-rich and stressful microenvironment that is likely to impose strong selective pressures arising from both host-mediated filtering and microbial interactions. In this highly selective context, only a subset of microbial taxa may possess adaptive traits, such as rapid nutrient uptake (Torres-Cortés et al., 2018), detoxification of reactive oxygen species (Gerna et al., 2020), or the ability to form protective structures like biofilms. This environment might trigger microbial competition, either through nutrient exploitation or via antagonistic interactions such as antibiosis (Russel et al., 2017, Garin et al., 2025). These competitive interactions may be influenced by metabolic resource overlap or by phylogenetic proximity between strains, as proposed by the competition-relatedness hypothesis, which suggests that closely related microorganisms are more likely to compete intensely (Russel et al., 2017; Machado et al., 2021). However, this hypothesis has rarely been tested in the context of plant microbiota, and never, to our knowledge, during the seed-to-seedling transition.

In this study, we employed a SynCom approach on common bean (*Phaseolus vulgaris*) seeds to investigate how bacterial traits and community composition influence strain colonization success during the seed-to-seedling transition. A collection of 30 strains isolated from common bean seeds was characterized based on their phenotypic traits (metabolic profiles and growth dynamics), and their genomes were sequenced to identify relevant functional traits (Arnault et al., 2024). Using these strains, SynComs were designed using in silico simulations of all possible community compositions, allowing us to select a subset of 30 SynComs spanning a wide range of phylogenetic diversity while maintaining a constant community size. This rational approach enabled us to test how phylogenetic proximity influences community assembly outcomes under controlled conditions. Based on the competition-relatedness hypothesis, we predicted that individual strains would exhibit higher transmission success from seed-to-seedling when phylogenetically distant from the other members of the community. To disentangle the relative contributions of host selection and microbial interactions to seedling colonization outcomes, each strain was also inoculated individually under the same experimental conditions. By addressing these, we aim not only to deepen our understanding of seed-to-seedling microbiota transmission but also to provide a framework for anticipating and potentially manipulating these processes to enhance microbiome engineering success (Russ et al., 2023; Joubert et al., 2025).

### Material and methods

### Strain selection and SynCom design

To build the SynComs, thirty bacterial strains were selected from a culture collection of 1,276 strains isolated from 8 different common bean genotypes (Flavert, Linex, Facila, Contender, Vanilla, Deezer, Vezer, and Caprice). Strains were selected according to (i) their abundance and prevalence of their *gyrB* sequences in seed and seedling communities and (ii) their phylogenetic diversity (Arnault et al., 2024). SynComs of 8 strains were designed as it represented the median observed richness of common bean seed microbiota (Chesneau et al., 2022). All combinations of 8 out of 30 strains (5,852,925 combinations) were simulated. A phylogenetic tree of the 30 strains was constructed using automlst (commit b116031) with default parameters and 1,000 bootstrap replicates (Alanjary et al., 2019). Using the simulated compositions and the phylogenetic tree, Faith’s phylogenetic diversity of each SynCom was calculated using the *pd* function of the picante package v1.8.2 (Kembel et al., 2010). The phylogenetic diversity across all the simulated SynComs was normally distributed, with a minimum of 1.6, maximum of 5.0 and mean of 3.7 (FigS1). Five groups of phylogenetic diversity corresponding to the following quantiles: 5%, 25%, 50%, 75% and 90% (FigS1) were selected. Two independent experiments including 15 SynComs each were performed. For each experiment, we selected three SynComs per phylogenetic group. We made sure that these three SynComs did not share any strain between them (using the homemade script *selectTriplets.pl*). For the second experiment, we selected SynComs using the same method and by ensuring that each SynCom shared a maximum number of 4 strains with SynComs from the first experiment. To sum up, 30 SynComs (6 SynComs per phylogenetic group x 5 phylogenetic groups) were designed (Fig1).

**Figure 1:**
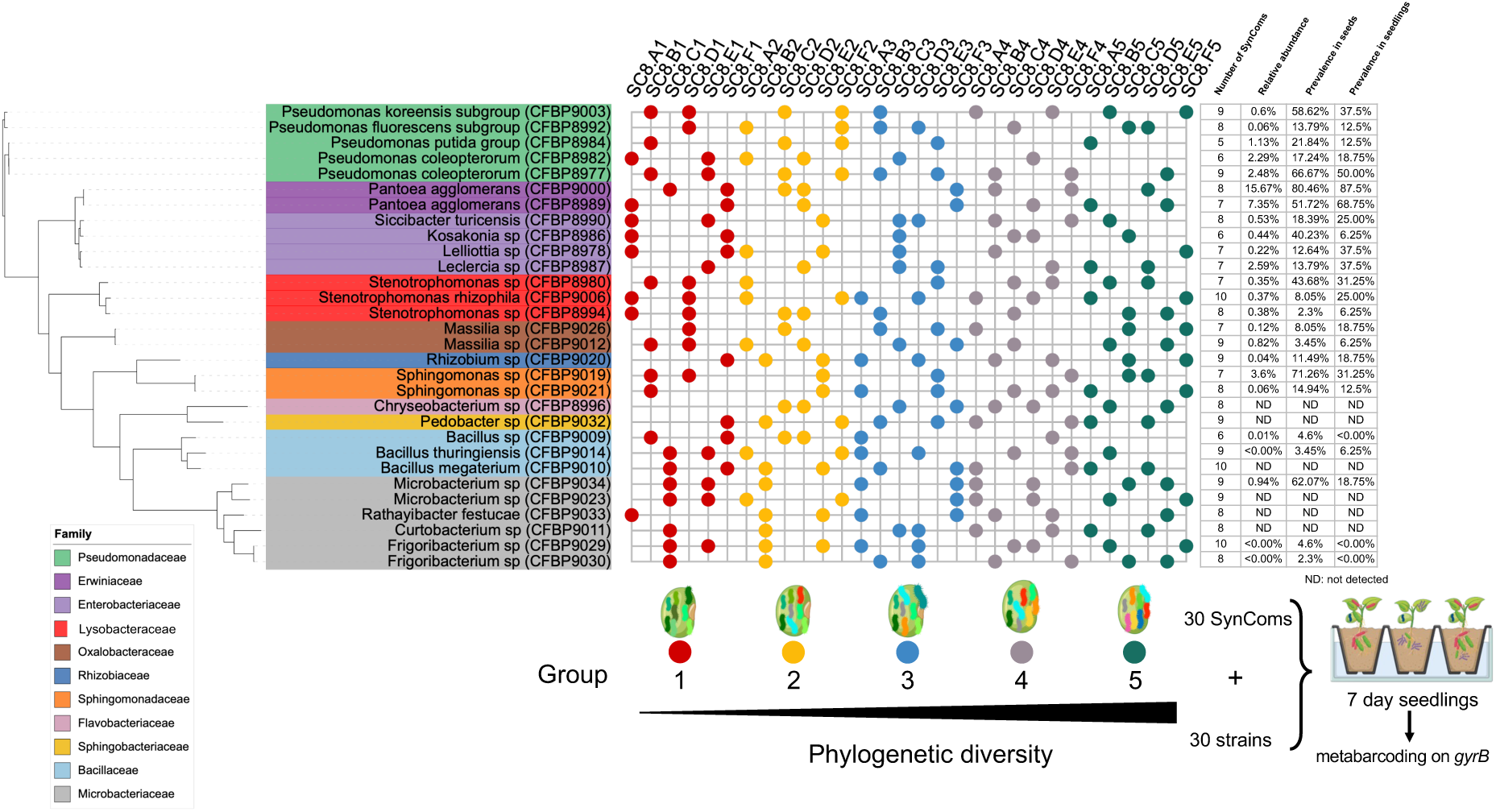
Overview of the experimental design, strain diversity and SynCom compositions. Thirty seed-borne bacterial strains isolated from common bean were employed in this work. Each strain has a different *gyrB* amplicon sequence variant (ASV) that can be distinguished from the others by sequencing. The relative abundance and prevalence of each *gyrB* ASV in seeds and seedlings are summarized on the dataframe (right). A total of 30 SynComs of 8 strains with contrasted phylogenetic diversity were constructed and seed-inoculated. All 30 strains were also individually seed-inoculated. The relative abundance of each strain in seedling was measured through *gyrB* amplicon sequencing.

### Strain phenotyping

To gain insights on potential microbial traits related to colonization efficiency, the selected strains were phenotyped for their growth and metabolic capacities. For growth capacity, bacteria were first cultivated on TSA media during 2 days at 18°C. Three isolated colonies were picked up and cultured overnight in TSB10 at 18°C under constant agitation (180 rpm). Cultures were centrifuged (4,000 g during 10 min) and pellets were resuspended in 10mM MgSO_4_ at an OD_600nm_ of 0.25. Twenty µl of bacterial suspensions were inoculated in triplicate on a microplate containing 180 μL of TSB 10%. Plates were incubated at 25°C during 48h under constant agitation (500 rpm). The package gcplyr v1.12.0 (Sprouffske et al., 2016) was used to estimate the different growth parameters (e.g. growth rate, carrying capacity, see Table S1).

Metabolic capacities of each strain were characterized with Biolog GEN III MicroPlate. Bacteria were cultivated on TSA during 2 days at 18°C. Single colonies were resuspended in inoculating fluid (IF-A) at a cell density of 92% transmittance. GEN III MicroPlates were incubated at 18°C for 3 days into the Omnilog Phenotype Microarray System. Microplates were monitored every 15 minutes. Three independent biological replicates were performed for each strain. For Biolog, data were exported as csv and analyzed with the Micro4Food PM pipeline (Acin-Albiac et al., 2020). Briefly, blanks were subtracted in each plate using the function PM.BgCorrectionMRT. Active and non-active profiles were separated according to the method developed by Vehkala et al., 2015 with an activity threshold set up to 50. Normalization between GEN III plates was performed after fitting a base active curve (Vehkala et al., 2015). Metabolic parameters were computed with the free spline method of the grofit package (Kahm et al., 2010). Substrates were categorized as active (1) or non-active (0). For some analyses, only wells corresponding to carbon source found on common bean seed exudates were kept (FigS5). This binary dataset was used to output pairwise metabolic distance between strains using the *mFD* package v1.0.7. Then, per strain and SynCom, the mean metabolic distance was obtained by calculating the mean metabolic distance between the 7 pairs of strains.

### Seed exudates profiling

To evaluate the metabolic capacity of strains on seeds, seed exudates were collected from bean seeds using 3 subsamples of 100 grams of seed (approximately 500 seeds for each subsample). Seeds were rinsed three times with sterile water and soaked in 250 mL of sterile distilled water overnight at 4°C under constant agitation (150 rpm). Solution was filtered through 0.2-μm-pore-size filters and lyophilized. The resulting powder was weighted and sent for GC-MS analysis (P2M2 facility). Based on seed exudates composition (Table S2), 17 substrates present in GEN III plates were selected. These selected substrates were used to calculate metabolic distances between strains (Exudates_distance).

### Strain et SynCom inoculation

All experiments were performed using a commercial seed lot of Flavert genotype from Vilmorin-Mikado (France). Since seed surface sterilization is not improving the inoculation success of SynComs (Arnault et al., 2024), seeds were only rinsed three times using sterilized water before inoculation. Each bacterial strain was grown on TSA10 for 2 days and resuspended in sterilized water. Each strain was adjusted at a population size of 10^7^ CFU/mL according to OD_600nm_. SynComs were prepared by mixing the same volume of each bacterial suspension. The concentrations of each bacterial suspension and SynCom were verified by dilution and plating on TSA10. Number of CFU was recorded 4 days after incubation at 18°C (FigS2). Seeds were inoculated with 2 mL of bacterial suspensions per gram of seed for 30 min under constant agitation (70 rpm) at 18°C. Excess inocula were removed using a sterile strainer, and seeds were dried for 30 min under a sterile laminar flow. Seed bacterial community biomass was measured after inoculation on eight seeds per condition. To do so, each seed was soaked in 2 mL of sterile water at 4°C under constant agitation (220 rpm) overnight. The resulting suspensions were then plated on TSA10 and microbiota profiling was performed on the same eight suspensions.

### Plant growth and DNA extraction

Inoculated seeds (n=40 per SynCom, n=120 per individual strain) were sown in a non-sterile potting soil (Traysubstrat Klasmann-Deilmann France) in a growth chamber for SynComs and greenhouse for individual strains (16 h day at 23°C, 8 h night at 20°C, 70% humidity). After 7 days of growth (BBCH stage 12, two full leaves unfolded), seedling roots were cleaned of potting soil excess by hand shaking and using sterilized water. Whole seedling samples (n=8 per SynCom, n=6 per individual strain) were first crushed with a roller. Then 2 mL of sterilized water was added, and the samples were ground for 30 s using a stomacher. DNA was extracted using 200 μL of the crushed suspension with the NucleoSpin® 96 Food kit (Macherey-Nagel, Düren, Germany) following the manufacturer’s instructions.

### Metabarcoding on *gyrB* gene and taxonomic classification

To assess the microbial community compositions, the following metabarcoding approach was performed on the inocula, inoculated seeds and seedlings of the different SynComs. For inocula, 200 μL of each fresh inoculum was instantly stored at -80°C before DNA extraction. For inoculated seeds, individual seeds were soaked as described before for CFU counting and 200 μL of each suspension obtained was stored at -80°C before DNA extraction and suspension.

To ensure accurate strain traceability, *gyrB* amplicon sequencing was performed across inocula, seeds, and seedlings samples. The first PCR was performed with the primers gyrB_aF64/gyrB_aR353 (Barret et al., 2015). PCR reactions were performed with a high-fidelity Taq DNA polymerase (AccuPrimeTM Taq DNA polymerase Polymerase System, Invitrogen, Carlsbad, California, USA) using 5 µL of 10X Buffer, 1 µL of forward and reverse primers (100 µM), 0.2 µL of Taq and 5 µl of DNA. PCR cycling conditions were done with an initial denaturation step at 94°C for 3 min, followed by 35 cycles of amplification at 94°C (30 s), 55°C (45 s) and 68°C (90 s), and a final elongation at 68°C for 10 min. Amplicons were purified with magnetic beads (Sera-MagTM, Merck, Kenilworth, New Jersey). The second PCR was conducted to incorporate Illumina adapters and barcodes. The PCR cycling conditions were: denaturation at 94°C (2 min), 12 cycles at 94°C (1 min), 55°C (1 min) and 68°C (1 min), and a final elongation at 68°C for 10 min. Amplicons were purified with magnetic beads and pooled. Concentration of the pool was measured with quantitative PCR (KAPA Library Quantification Kit, Roche, Basel, Switzerland). Amplicon libraries were mixed with 10% PhiX and sequenced with two MiSeq reagent kits v2 500 cycles (Illumina, San Diego, California, USA). A blank extraction kit control, a PCR-negative control and PCR-positive control (*Lactococcus piscium* DSM6634, a fish pathogen that is not plant-associated) were included in each PCR plate.

The bioinformatic processing of the amplicons was performed in R. In brief, primer sequences were removed with cutadapt 2.7 (Martin, 2011) and trimmed fastq files were processed with DADA2 version 1.22.0 (Callahan et al., 2016). Chimeric sequences were identified and removed with the removeBimeraDenovo function of DADA2. Amplicon Sequence Variant (ASV) taxonomic affiliations were performed with a naive Bayesian classifier (Wang et al., 2007) with our in-house *gyrB* gene database (Briand et al., 2025). We identified several Single Nucleotide Polymorphisms (SNPs) in our inocula sample that were artificially increasing sample richness. Thus, the post-clustering algorithm LULU v0.1.0 (Froslev et al., 2017) was used to merge *gyrB* ASV at an identity threshold of 98%. Unassigned sequences at the phylum level, *parE* sequences (a *gyrB* paralog) and singletons were filtered. Only ASVs sharing 100% sequence identity with the inoculated strains were considered to be part of the initial SynComs.

### Statistical analyses

#### Microbiota profiling and statistical analyses

Microbial community analyses were conducted using the Phyloseq package v1.44.0 (McMurdie and Holmes 2013). Figures were generated using ggplot2 v3.4.3. Dataset from the two different experiments were merged and the final data set was rarefied (FigS3) using a coverage-based method proposed by Chao and Jost (2012) using the phyloseq_coverage_raref function from metagMisc package v0.5.0 (Mikryukov and Mahé, 2024; Chao and Jost, 2012). Beta diversity analyses were made using BrayCurtis distance, PCoA ordination and permutational multivariate analysis of variance [adonis2 function of vegan v2.6.4 (Oksanen et al., 2019), 999 permutations]. Wilcoxon tests were used to compare colonization success of SynComs. Linear regressions were obtained using *lm* function from *stats* package and assumptions of normality of the residuals and homoscedasticity were checked by plotting the models using the *plot* function in R. Linear mixed model was obtained using *lmer* function from *lme4* package v.1.1.37.

#### Genome assembly and analyses

To gain insights on potentially relevant genomic traits involved in colonization competitiveness, a genomic analysis of the 30 strains was conducted. Genomes of the strains were sequenced at BGI (China) using DNBSEQ technology and assembled with Spades v3.15.3 using the default k-mer parameters (-k 21, 33, 55, 77, 99, and 127) and the following options: –cov-cutoff auto, –isolate (Prjibelski et al., 2020). Genomes are available on NCBI using the BioProject ID PRJNA1041598. Genome information of the different strains obtained are summarized in Table S3. Functional annotations were carried out with prokka v1.14.6 (Seemann 2014) and orthology assignment was performed with DIAMOND v2.1.3 (Buchfink et al., 2021) on the eggNOG 5.0 database (Huerta-Cepas et al., 2019). Orthologous groups (OGs) associated with high / low seedling colonization capacities were identified with Scoary for SynCom contexts using our own categorization (Brynildsrud et al., 2016) and Scoary2 for individual contexts (Roder et al., 2024) using a presence/absence table of OGs. From the genomes, a matrix of traits that include resource acquisition, resource use, life history traits and stress tolerance traits was obtained using the *microTrait* package v1.0.0 (Karaoz et al., 2022). The granularity 3 matrix obtained was used to classify the strains in 6 different clusters using a hierarchical clustering approach (see FigS4 for details).

For each strain in each SynCom, the mean phylogenetic distance from other strains within the SynCom was calculated. To do so, a phylogenetic tree was constructed using automlst (commit b116031) with default parameters and 1000 bootstrap replicates (Alanjary et al., 2019). Then, cophenetic distance between pairs of strains was calculated using the *cophenetic()* function in R. Then, mean phylogenetic distance per strain in each SynCom was obtained by calculating the mean between the 7 phylogenetic distances.

#### Random forest analysis

We used a custom python-based script to implement a random forest approach to predict bacterial strain colonization capacity using a regression model. Specifically, we aimed to predict the exact log-fold change in abundance between the seed and the seedling for each strain (log2FC). The log2FC was computed as follows: [log2FC: log2(mean relative abundance in seedling of SynCom_x_ (%) / mean relative abundance in seed of SynCom_x_ (%))]. First, a dataset gathering all strain descriptors (features) assessed in this study was built (232 observations described by 16 different features, see Table S4 for details). Then, the dataset was randomly split in a train set (80%, n = 186) and a test set (20%, n = 46). The trainset was used to train the model to predict the desired known log2FC output values. The test set, which contains data that was not seen by the model during training, was used to assess the accuracy of the predictions.

For the regression model, the ‘*RandomForestRegressor*’ package from ‘*sklearn.ensemble*’ was used (Pedregosa et al., 2011) (1000 trees, Friedman Mean Square Error impurity improvement criterion). To evaluate the performance of the regression model, we compared our predicted log2FC values to the actual observed values in the test set, using a linear model regression. To understand which features were the most contributing in the regression prediction, we used the ‘feature_permutation’ method to assess the importance of each feature on the accuracy of the model (100 permutations).

## Results

### Effective modulation of seed and seedling microbiota phylogenetic diversity using SynComs

Thirty SynComs of eight strains were inoculated on seeds (Fig1). These SynComs present distinct levels of phylogenetic diversity (*Material and Methods*). Seed (Fig2-A) and seedling (Fig2-B) microbiota composition were significantly modified after SynCom inoculation with 89% and 79% of explained variance, respectively (p < 0.001). The colonization success of seedling per SynCom was estimated by cumulating the relative abundance of *gyrB* ASVs of each strain. This cumulative relative abundance ranged from 9.1% to 94.4% with an average of 77.8% and was significantly different from uninoculated seedlings (Fig2-C; Wilcoxon test; p-value < 0.05). In order to limit the influence of non-inoculated bacteria in our subsequent analyses, we decided to exclude seedling samples with less than 50% cumulative relative abundance (30 out of 240).

**Figure 2:**
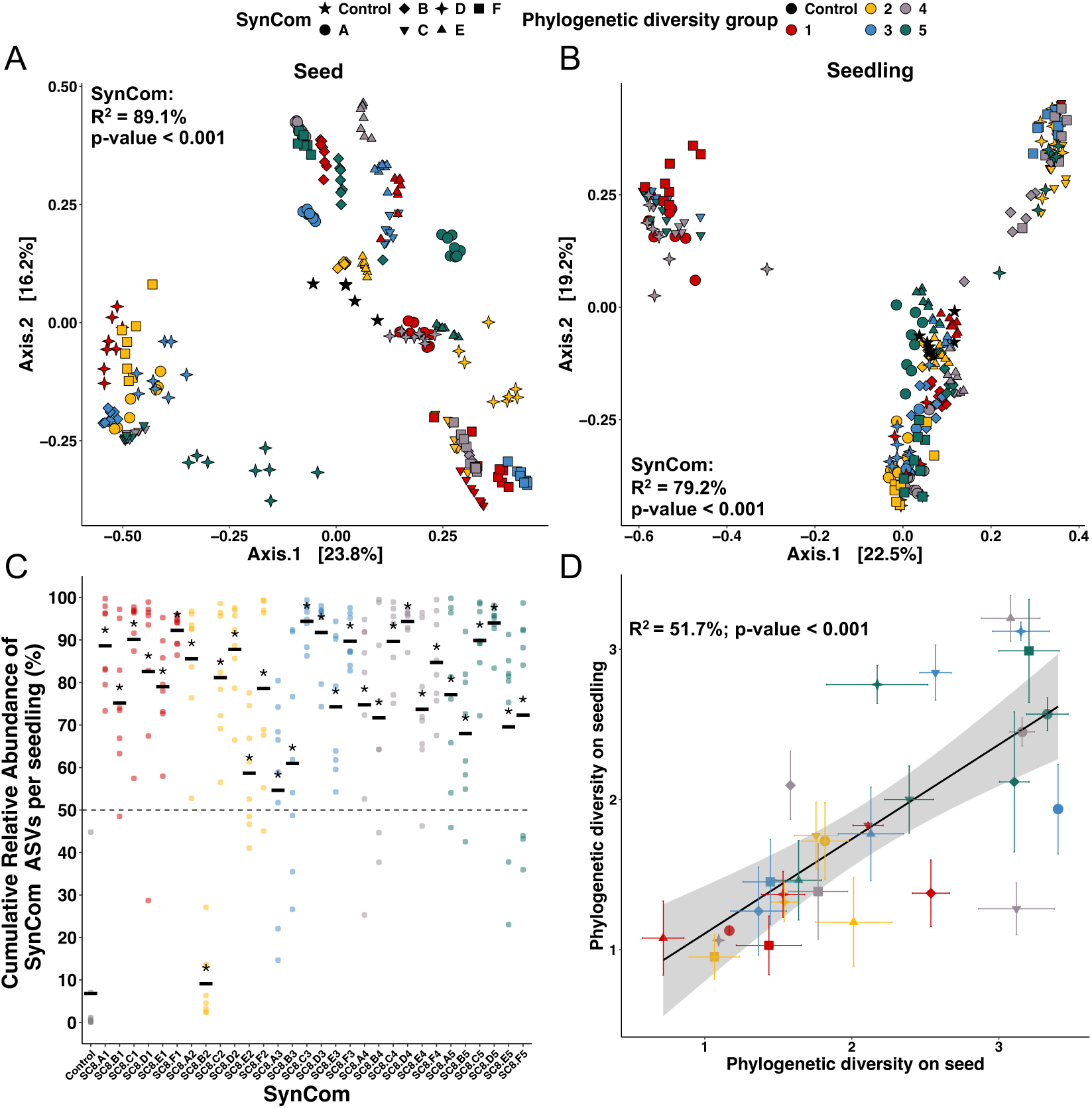
Effect of SynCom inoculation on seed and seedling bacterial communities. (A-B): Influence of SynCom inoculation on seed (A) and seedling (B) bacterial community compositions. PCoA ordinations were derived from Bray–Curtis distances (PERMANOVA test; SynCom effect: R^2^ = 89.1% for seed and R^2^ = 79.2% for seedling, p-values < 0.001). Each point represents a single sample (seed or seedling) and is colored by its phylogenetic group. A unique combination of shapes (A-F, 6 replicates) and color (group of diversity) is displayed per SynCom. (C) : Cumulative relative abundance of SynCom ASVs per seedling (%). Asterisks indicate significant increase of the cumulative relative abundance of SynCom ASVs compared to uninoculated control (Wilcoxon test, p-values < 0.05). Each point represents one seedling and is colored based on phylogenetic diversity. The mean cumulative relative abundance of SynCom ASVs are represented by the black line. The dashed line indicates the relative abundance of 50% and allows to visualize the 30 samples that were removed from the dataset for further analysis of seedling microbiota. (D) : Positive linear regression between the phylogenetic diversity of SynCom on seedling and its phylogenetic diversity on seed (R^2^ = 51.7%, p-value < 0.001).

The theoretical phylogenetic diversity of the SynComs was positively correlated with the observed phylogenetic diversity of the inocula (FigS6A; Linear regression; R^2^ = 60.3%; p-value < 0.001), seed microbiota (FigS6B; R^2^ = 21.9%; p-value = 0.006) and seedling microbiota (FigS6C; R^2^ = 25.8%; p-value = 0.003). Finally, SynComs inoculation did alter the seed and seedling microbiota, as the observed phylogenetic diversity of seed was positively correlated with the phylogenetic diversity of seedlings (Fig2-D; R^2^ = 51.7%; p-value < 0.001).

### Variable strain colonization success from seed-to-seedling

Taxonomic profiles revealed that some strains did not successfully transmit from seeds to seedlings, while some presented high seedling colonization (FigS7). Seedling colonization capacity of each strain in each SynCom was assessed by using a log2FoldChange (log2FC): log2(%Relative abundance in seedling/%Relative abundance in seed). Globally, strains displayed more often low rather than high seedling colonization capacities (Fig3). The strains exhibited varying seedling colonization capacities, with a minimum mean log2FC of -4.7 for a strain belonging to the *Pseudomonas putida group* (CFBP8984) and a maximum mean log2FC of 2.6 for a strain of *Stenotrophomonas* sp. (CFBP8994) (Fig3). Most strains showed high variability in seedling colonization capacity according to the SynCom composition (Fig3). For instance, *Pseudomonas coleopterorum* (CFBP8977) had a mean log2FC of -1.7, ranging from -4.8 in SynCom8.B1 to 0.9 in SynCom8.E1. Only one *Kosakonia* sp. strain (CFBP8986) had a positive and stable log2FC across all SynComs (from 2.2 to 2.4).

**Figure 3:**
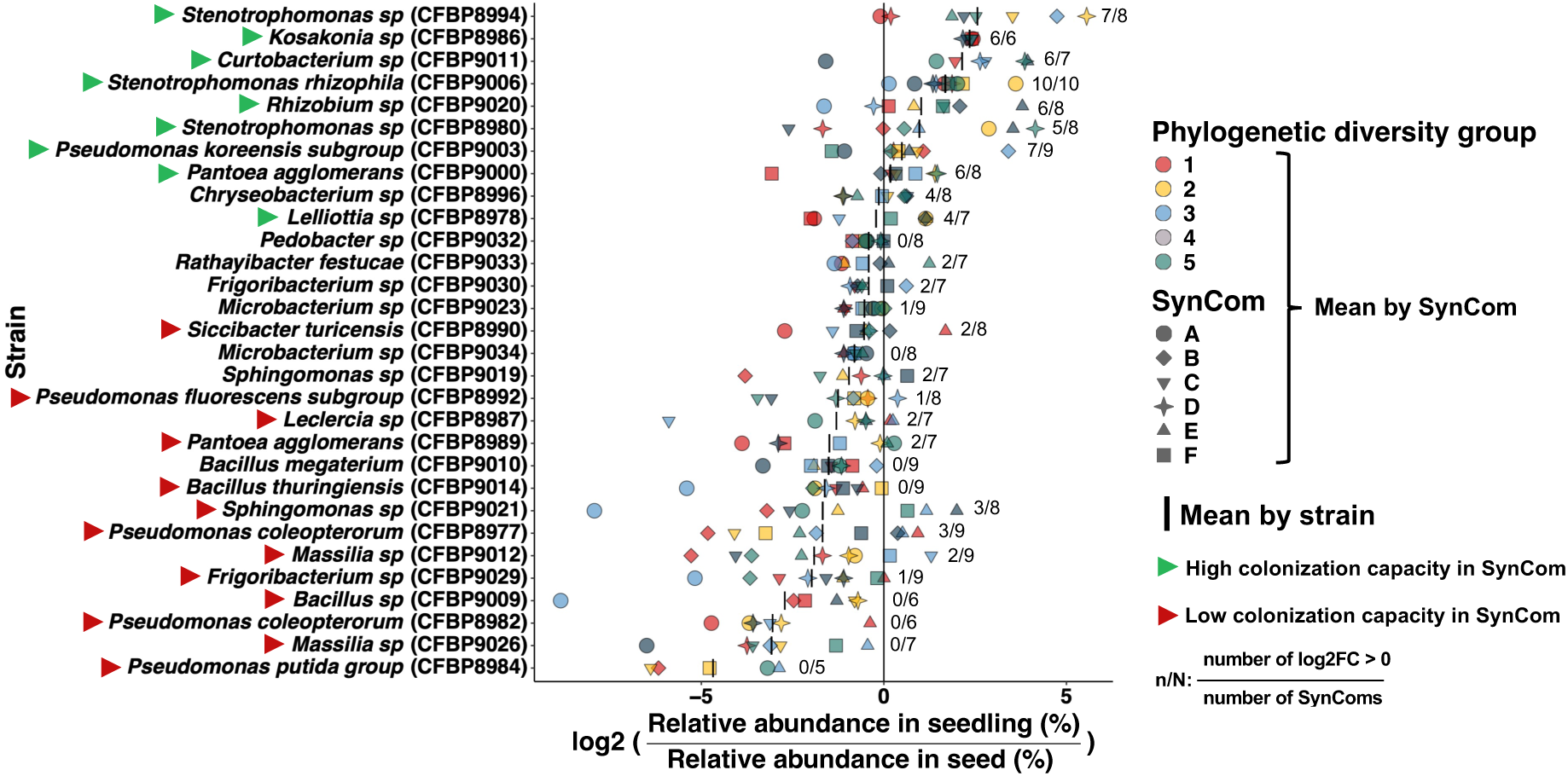
Seedling colonization capacity of each strain within each SynCom. Strain’s ability to colonize seedlings compared to their initial relative abundance on seeds, depending on the SynCom composition. The following ratio, hereafter called log2FC, was calculated to assess this trait: log2(mean relative abundance in seedling of SynCom_x_ (%) / mean relative abundance in seed of SynCom_x_ (%). The log2FC is plotted for each SynCom (combination of color and shape) and the black line represents the mean across all SynComs in which the strain was included. For each strain, the number of SynCom for which log2FC ratio > 0 is indicated (n) as well as the number of SynComs in which the strain is included (N). If the ratio n/N > 50% and the strain relative abundance > 0.05% in at least seed or seedling, the strain is considered as a high colonizer in SynComs (green arrow) and if n/N < 50% as low colonizer in SynComs (red arrow). These high and low colonizer categories were used for the comparative genomic analysis later (Scoary). Strains are ranked on the y-axis according to their average log2FC across all SynComs (black line).

Based on these initial results, strains were categorized as high and low colonizers. To avoid biases coming from the detection limit, strains were not categorized if their relative abundances were under 0.05% in both seed and seedling. This led to a group of high colonizers composed of 9 strains (green arrows) and a group of low colonization capacity composed of 13 strains (red arrows; Fig3).

To decipher the relative importance of strain identity *versus* SynCom composition in driving the seedling colonization capacity, each strain was also seed-inoculated individually (Fig4 & FigS8). A strain was defined as transmitted if its relative abundance in seedling was superior to 0.05%. Individual strains showed highly contrasted transmission rate and relative abundance in seedlings (Fig4). For instance, *Microbacterium* sp. (CFBP9034) was never detected in seedlings, while a strain of the *Pseudomonas fluorescens* subgroup (CFBP8992) was always transmitted (Fig4 & FigS8). Across strains that were always transmitted, their relative abundances in seedlings were highly variable, ranging from 3.4% for *Curtobacterium* sp. (CFBP9021) to 93.5% in a strain of the *Pseudomonas fluorescens* subgroup (CFBP8992). The next step was to compare seedling colonization capacities of every strain when inoculated alone or in SynComs. The seventeen strains that were transmitted to seedlings in both contexts (i.e. individual inoculation and SynComs) were kept for further analyses. Interestingly, strain colonization in SynComs could not be predicted from their individual profiles, as shown by the different results obtained either in individual inoculation (Fig4) and in SynComs (Fig3), being consistent with the lack of correlation between their relative abundances in these two contexts (FigS9). Prominent examples are strains of the *P. fluorescens* subgroup (CFBP8992), *Siccibacter turicensis* (CFBP8990) and *Leclercia* sp. (CFBP8987) that were respectively ranked 1^st^, 5^th^ and 6^th^ in individual strain inoculation experiments, while they presented low colonization capacity in SynComs (Fig3 and FigS9).

**Figure 4:**
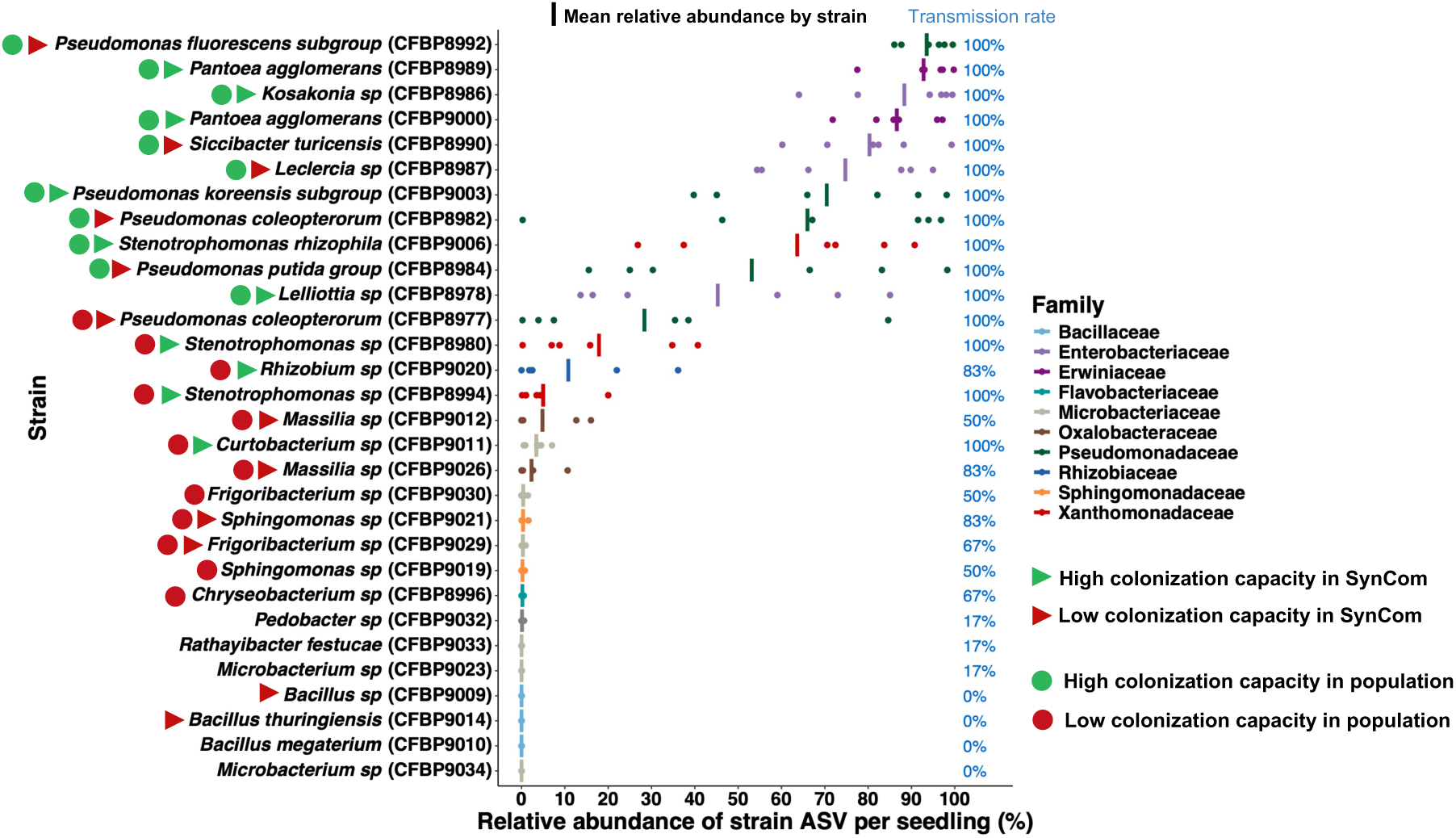
Seedling colonization capacity of each strain inoculated individually. Relative abundance of strain ASV per seedling (%) when inoculated individually on seed. Each point represents the relative abundance in one seedling (n=6) and the mean relative abundance per strain is indicated with a vertical dash. The transmission rate (%) of each strain is indicated in blue on the right of the plot. A strain was considered as transmitted if its relative abundance > 0.05% on a given seedling. Based on the mean relative abundance, strains were categorized as high and low seedling colonizers in individual contexts using Scoary2. These high and low categories were used for the comparative genomic analysis later (Scoary2). High (green) and low (red) colonization capacity in SynComs are also reported (arrow).

### Strain identity, phylogenetic distance et metabolic capacities are the main predictors of seedling colonization success

To elucidate which strain- and community-level traits could explain the variable transmission of strains from seed-to-seedling, a random forest regression model was built to attempt predicting the log2FC of each strain and identify the key features involved. For this purpose, several traits were acquired (see Table S4 for a summary), including (i) genome-based functional profiling (FigS4), (ii) strain growth and metabolic capacities (TableS1 & FigS5) and (iii) strains phylogenetic and metabolic relatedness within each community.

Results of the random forest model are presented in Figure 5. We obtained an accuracy score of 47% (panel A) in attempting to predict the exact log2FC value of a given strain, with a Friedman mean square error of 1.38. We could fit a linear model between our predicted and observed values of the test set (y = 1.03x + 0.28), with an adjusted R^2^ of 0.48 (P < 0.001, Panel B). We observed that our model had an overall efficient prediction proportionality, with a slope very close to 1. However, the model tended to overestimate the actual log2FC by +0.28 (intercept). The strain features with significant impact on the accuracy of the model are shown in Panel A. The most important features for accurate prediction of strain colonization capacity in terms of R^2^ contributions were first their identity (“Strain” R^2^: 0.3961), followed by the mean phylogenetic distance between all members of the SynCom where they belonged (“Phylogenetic_distance”: 0.1185), their metabolic distance with other strains based on seed exudate metabolization (“Exudates_distance”: 0.1158), the SynCom (0.097) and their ranks in terms of number of seed exudates metabolized compared to the other strains in the SynCom (“BIOLOG_exudates”: 0.075). Other parameters contributed significantly as well, but for less than 0.05 accuracy.

**Figure 5:**
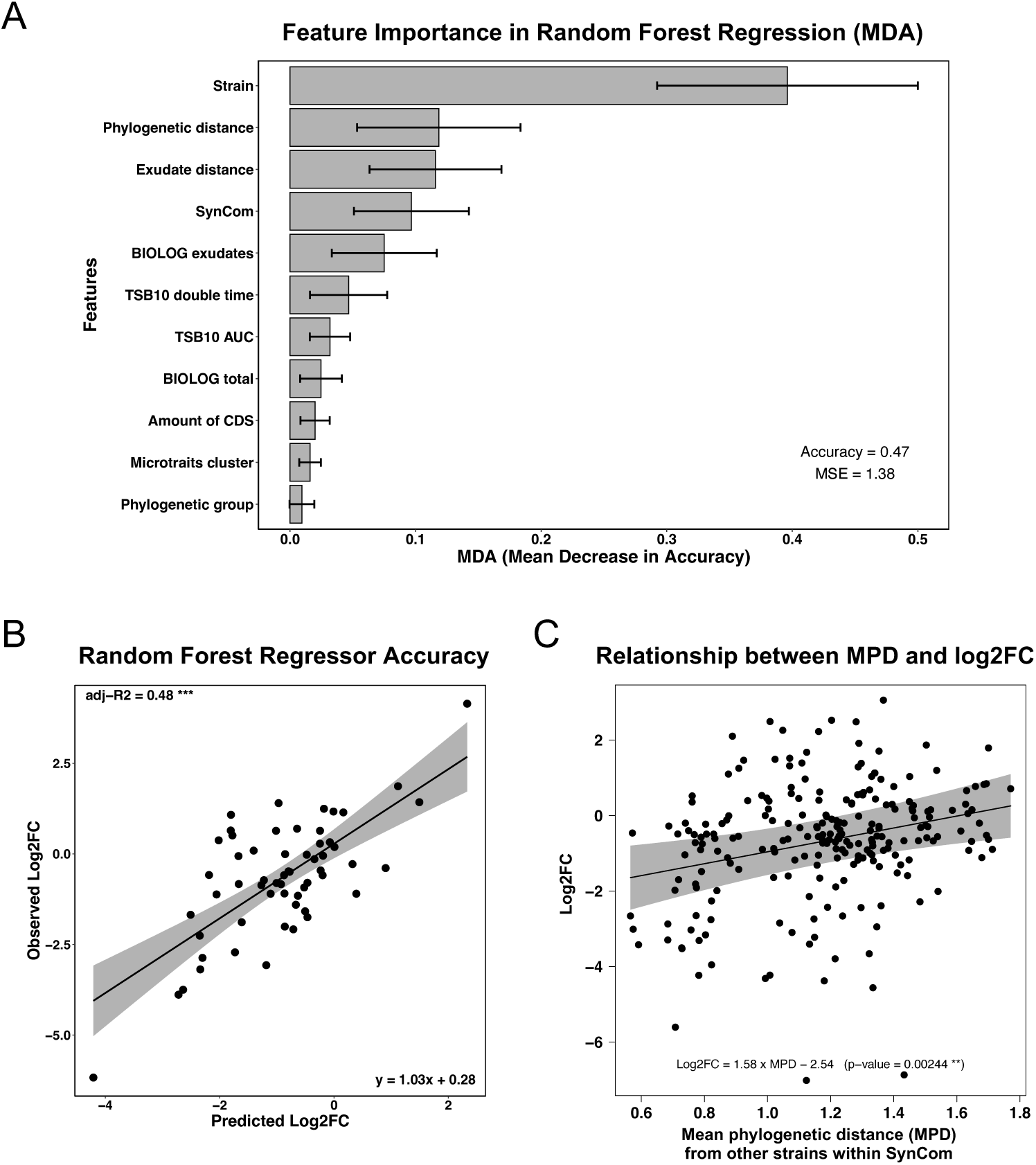
Results of the random forest analysis by regression. (A) Features importance of the Random Forest Regression model. Strain: Identity of the strain; Phylogenetic distance: the average phylogenetic distance of the strain with the others in the SynCom; Exudate distance: the average metabolic distance (exudates-like) of the strain with the others in the SynCom. SynCom: Identity of the SynCom. BIOLOG exudates: the rank of the strain in terms of usable exudate-like substrates in the SynCom; TSB 10 doubling time: the rank of the strain in terms of growth (doubling time) in the SynCom. TSB10_AUC: the rank of the strain in terms of growth (Area under the curve) in the SynCom. BIOLOG total: the rank of the strain in terms of usable total substrates in the SynCom. Amount of CDS: the rank of the strain in terms of number of CDS in the SynCom. Microtraits cluster: Cluster of the strain based on the microtraits matrix obtained from genomes. Phylogenetic group: Initial phylogenetic group of the SynCom. See Table S4 for details. (B) Accuracy of the model: linear regression between the observed log2FC and their predicted log2FC using the test set. (C) Linear mixed model between the log2FC and the mean phylogenetic distance (MPD) from other strains within SynCom. The log2FC from seed-to-seedling was calculated for each strain in each SynCom as well as its mean phylogenetic distance from other strains within the SynCom. A linear mixed model was made using the following formula: log2FC ∼ MPD + (1|strain). (MPD as fixed effect; strain: strain identity as a random effect).

Because the mean phylogenetic distance (MPD) was the second most important feature after strain identity (Fig5-A), we built a linear mixed model to explain the log2FC as a function of the mean phylogenetic distance with strain identity as random effect (Fig5-C). We found that log2FC was positively correlated with the mean phylogenetic distance from other strains, meaning that strains colonization is higher when they are more distant from the others within the SynCom (Log2FC = 1.414MPD – 3.708; p-value < 0.01). Linear regressions between log2FC and mean phylogenetic distance were also tested for each strain. Only *Lellotia* sp. (CFBP8978) and *Stenotrophomonas rhizophila* (CFBP9006) presented significant linear regressions (p-values < 0.05). Interestingly, while log2FC of *Lellotia sp* was positively correlated to the mean phylogenetic distance, *Stenotrophomonas rhizophila* was negatively associated (FigS10).

### Genetic traits of high *versus* low seedling colonization in individual and SynCom contexts

Because strain identity was the major feature contributing to the accuracy of the random forest analysis, we assessed whether genetic determinants could be involved in seedling colonization success by analyzing the genomes of our 30 strains. A first comparative genomic analysis was performed between the high and the low colonizers in SynCom contexts using Scoary. To highlight only the best ortholog group (OG) candidates, only OGs with a sensitivity and specificity higher than 75% were kept. This resulted in a list of 6 OGs associated with a high seedling colonization capacity: COG0376, COG4558, COG4559, COG2214, COG4229 and COG2269 (table 1). Three of these OGs were associated with inorganic ion transport and metabolism including a catalase/peroxidase coded by *katG* gene (COG0376) and two protein-coding genes involved in the transport of hemin (COG4558, COG4559).

**Table 1:** COGs specific to high colonization capacity in SynCom contexts.

| COGs | Sensitivity | Specificity | Function |
| --- | --- | --- | --- |
| COG0376 | 100.0 | 76.9 | Catalase (peroxidase I), KatG |
| COG2214 | 77.8 | 84.6 | Hypothetical protein |
| COG4558 | 88.9 | 76.9 | Heme import, HmuT |
| COG4559 | 88.9 | 76.9 | Heme import, HmuV |
| COG4229 | 77.8 | 76.9 | Enolase-phosphatase E1 |
| COG2269 | 77.8 | 76.9 | Elongation factor P--beta-lysine ligase, EmpA |

Then, a second comparative genomic analysis was performed between the high and the low colonizers in individual inoculation contexts. To do so, Scoary2 was used with the mean relative abundance as input values after removing strains with low detection (mean relative abundance < 0.05%). This gave a group of 10 strains with high seedling colonization capacity (green points on Fig4) and a group of 12 strains with low seedling colonization capacity in individual context (red points on Fig4). As before, only OGs with a sensitivity and specificity higher than 75% were kept. This resulted in a list of 95 OGs associated with a high seedling colonization capacity in individual context (Table S5). Most of these OGs had unknown functions (n=38), were involved in amino acid transport and metabolism (n=14), post-transcriptional modifications (n=5), cell wall (n=5) and stress resistance (n=5).

## Discussion

### Strain identity and community composition are the main drivers of seed-to-seedling microbiota assembly

By testing strain colonization capacity in both individual and SynCom contexts, we aimed to disentangle the relative contributions of microbial traits and interactions in shaping microbiota transmission from seed-to-seedling. By inoculating a series of phylogenetically contrasted SynComs, we were able to change the phylogenetic diversity and composition of both seed and seedling microbiota in a consistent manner. We observed that the best seedling colonizers in individual inoculation contexts were not necessarily those that performed best within SynComs. Thus, seedling colonization success in community context is not predictable based on single strain inoculation. A random forest model was used to identify the strain-level traits and community-level parameters that best predict strain colonization success during the seed-to-seedling transition. Strain identity emerged as the most important predictor, suggesting that strain-specific features - particularly metabolic and functional traits - play a central role in determining transmission capacity in community contexts. Given the importance of these traits, we further explored the genomic features potentially underlying strain-specific transmission capacities, aiming to identify the genetic determinants that may govern successful transmission from seed-to-seedling.

### Identifying the key strain traits associated with high seedling colonization capacities

Using two different comparative genomic analyses, we investigated the potential traits involved in the high colonization capacity in individual and SynCom contexts. The first comparative genomic analysis was made between high *versus* low seedling colonization capacity in SynCom contexts. Only 6 candidate genes were detected in the majority of strains belonging to the high colonizers. Among these genes, the catalase-peroxidase KatG is the only one to be detected in 100% of the strains transmitted. KatG, is a bifunctional catalase-peroxidase which is expressed under conditions of oxidative stress (Yuan et al., 2021). This enzyme is involved in the detoxification of hydrogen peroxide (H_2_O_2_) and breaks down harmful peroxides in a number of plant-associated bacteria including *Bradyrhizobium japonicum* (R. Panek et al., 2004), *Enterobacter* (Taghavi et al., 2010) and *Bacillus* (Singha et al., 2021). KatG catalyzes the dismutation of H_2_O_2_ through oxidation of heme iron and reduces peroxides using electron donors (Zamocky et al., 2008). Interestingly two protein-coding genes (HmuT and HmuV) involved in the biogenesis of an ABC-type heme transporter system also belong to the list of genes associated with high colonizing strains (Muraki et al., 2016). This heme transporter could not only facilitate the acquisition of iron but also reduce the oxidative damage of H_2_O_2_. Since H_2_O_2_ production is induced by seed microbiota during germination and seedling emergence (Gerna et al., 2020), tolerance to oxidative stress is likely to be an important feature of efficient seedling colonization (Joubert et al., 2024). Reactive oxygen species (ROS) such as H_2_O_2_ and peroxides are not only produced by seeds during germination process but also by bacteria during intense metabolic activity and under toxic metabolic exposition during intermicrobial competition.

The second comparative genomics analysis carried out on the colonization of individual strains identified numerous candidate genes (n=95). Surprisingly, none of the genes identified in the first comparative genomic analysis were shared, highlighting that the genetic traits required to colonize seedling when inoculated individually are different from those needed in community contexts. Thus, the genes involved in resistance to oxidative stress were not detected. This could be explained by the differential production of hydrogen peroxide content depending on strain identity (Gerna et al., 2019). In other words, in a SynCom context, it is more likely to achieve high production levels of H_2_O_2_ because the chance of having a strain that elicits this response is greater and/or because bacteria can produce more H_2_O_2_ themselves. It is difficult to identify a particularly enriched functional category in the candidate genes. Beyond genes of unknown functions (n=38), we can nevertheless note the presence of many genes involved in the metabolism and transport of amino acids (n=14), in particular arginine and branched-chain amino acids (Table S5). The role of amino acids in (i) chemotaxis and (ii) growth of bacterial strains within the spermosphere has already been documented (reviewed in Nelson et al., 2004). For instance, *Enterobacter cloacae* required specific amino acids to grow in spermosphere environments (Roberts et al., 1996). As the use of amino acids as a source of carbon and nitrogen is not restricted to good colonizing strains, it is very likely that other mechanisms are involved in competence for this habitat. For instance, the two most specific OGs (COG3098 and COG3539) were potentially involved in biofilm formation and cell adhesion (Beloin et al., 2004; Kim et al, 2009). These properties might enhance seedling colonization, as biofilm formation and cell adhesion are known to favor bacterial colonization of the plant (Danhorn et al., 2007; Li et al., 2024, Mazo-Monzalzo et al., 2024).

Together, these results support the idea that distinct selective pressures act on strains depending on whether they colonize the seedling alone or within a microbial community. The identification of catalase-peroxidase KatG and heme transporter genes in high SynCom colonizers validates the adaptive prediction that tolerance to oxidative stress enhances survival and transmission through the seed-to-seedling transition. In contrast, candidate genes in individual inoculations point toward alternative adaptive routes—such as nutrient acquisition or adhesion—highlighting context-specific trait requirements. The complete lack of overlap between the two genomic profiles underscores the critical role of bacterial interactions in shaping colonization outcomes. These findings call for functional validation, for instance through targeted gene knockouts or high-throughput fitness profiling approaches such as RB-TnSeq, to confirm the causal role of these candidate genes in seedling colonization across different ecological contexts.

### Phylogenetic and metabolic relatedness is a predictor of seed-to-seedling transmission

Phylogenetic relatedness between strains emerged as the second most important predictor in the regression-based random forest model. We initially predicted that higher phylogenetic distance between SynCom members should increase the colonization success of strains belonging to that SynCom, as stated by the competition-relatedness hypothesis (Russel et al., 2017, Machado et al., 2021). We found a weak but positive correlation between a strain’s colonization success and its mean phylogenetic distance from other strains within SynCom. This suggests that reduced phylogenetic similarity—and potentially reduced niche overlap—may alleviate competitive pressures, thereby favoring successful colonization (Russel et al., 2017; Joubert et al., 2024). However, at strain level, the seedling colonization was positively correlated with phylogenetic distance for *Lelliottia sp* (CFBP8978) while it was negatively correlated for *Stenotrophomonas rhizophil*a (CFBP9006) suggesting that microbial interactions are also strain-specific and SynCom-dependent. This finding is crucial as it implies that the assembly of seedling microbiota cannot be accurately predicted solely based on individual strain capacity nor phylogenetic relationships between strains but actually requires a priori knowledge on the actual causal biotic interactions between microbial members. For instance, *Stenotrophomonas rhizophil*a (CFBP9006) showed better colonization success when it was exposed to closely related strains. This observation could be explained by an interference interaction through the Type VI Secretion System (T6SS). Indeed, a seed-to-seedling study on radish revealed that interference competition mediated through T6SS-contact dependent mechanisms was more prevalent among closely related strains (Garin et al., 2025). In our case, *Stenotrophomonas rhizophil*a (CFBP9006) has a complete T6SS but further investigation would be needed to confirm this type of microbial interaction.

During germination, seeds release many exudates that can serve as substrates for many microorganisms from the seed or the soil (Saccaram et al., 2025). This influx of resources creates new niches that can be exploited by metabolically adapted microorganisms, promoting their establishment in seedlings (Joubert et al., 2024). Supporting this, several metabolic and growth traits ranked among the top predictors of colonization success in our model. Specifically, high colonizers were enriched in traits related to amino acid metabolism and iron acquisition (e.g., heme utilization). This highlights the role of trophic versatility and metabolic adaptation in enabling strains to persist through the seed-to-seedling transition. However, we found no linear relationship between a strain’s colonization success and its mean metabolic distance to other SynCom members. This suggests that, unlike phylogenetic proximity, trophic similarity does not necessarily intensify competitive exclusion in this context. This lack of strong signal for metabolic competition may reflect the diversity and abundance of accessible resources during early plant development (Saccaram et al., 2025). Most strains in our SynComs were able to grow on a broad range of seed-derived substrates, reducing the likelihood of direct resource competition. Moreover, spatial heterogeneity within the spermosphere and developing seedling tissues could limit niche overlap at the microscale, further dampening the effects of metabolic competition (Schlechter et al., 2023). Altogether, these results suggest that while trophic competence is a prerequisite for colonization, its influence is modulated by ecological context and spatial structure, making it less predictive than phylogenetic relationships or oxidative stress-related traits in explaining colonization outcomes.

## Conclusion

Our results reveal that a phylogenetically diverse set of strains can achieve high seedling colonization success via distinct ecological and functional strategies. These strategies include divergent growth dynamics, contrasting metabolic profiles, and the presence of specific candidate genes linked to seedling competence. Notably, the success of seed-to-seedling transmission was highly context-dependent, varying significantly between single-strain inoculations and SynComs. This indicates that seedling colonization cannot be predicted solely based on individual strain behavior, emphasizing the influence of microbial interactions during community assembly.

From an applied perspective, these insights have strong implications for microbiota engineering. Designing effective SynComs for seed inoculation will likely require integrating both high-performing individual strains and complementary trait combinations that ensure robustness across varying community compositions. Our results suggest that screening strains solely in isolation may be insufficient to identify optimal candidates. Instead, a combinatorial approach that explores strain performance across diverse biotic contexts is needed. Yet, even with detailed knowledge of functional traits and phylogenetic structure, emergent properties from microbial interactions may limit the predictability of colonization outcomes. This underlines the need for predictive models that incorporate not only microbial traits, but also their interaction networks and community dynamics. Ultimately, advancing microbiota-based plant engineering will depend on our capacity to navigate this functional and interactional complexity—by designing microbial consortia that are both trait-diverse and interaction-compatible, and by validating their robustness under realistic ecological conditions.

## Supporting information

Supplementary tables Arnault 2026

## Acknowledgements

This research was conducted through the OSMOSE project (2020– 2022) in the framework of the regional programme ‘Objectif Végétal, Research, Education and Innovation in Pays de la Loire’, supported by the French Region Pays de la Loire, Angers Loire Métropole and the European Regional Development Fund. This work was also part of the 3rd Programme for Future Investments (France2030) and operated by the SUCSEED project (ANR-20-PCPA-0009) funded by the ‘Growing and Protecting crops Differently’ French Priority Research Program (PPR-CPA), part of the national investment plan operated by the French National Research Agency (ANR). Gontran Arnault PhD was supported by both INRAE (Plant Health and Environment division, HoloFlux Metaprogram) and the Région des Pays de la Loire.

We thank Muriel Bahut (ANAN platform, SFR Quasav) for amplicon sequencing and the Phenotic platform for the plant cultivation in greenhouse and climatic chambers (DOI: 10.17180/ykbz-2v85). We also acknowledge the contribution of the CIRM-CFBP (International Centre of Microbial Resource (CIRM) - French Collection for Plant-associated Bacteria, INRAE) (https://doi.org/10.15454/E8XX-4Z18) for strain preservation. We thank Théophile Trunck for his help in creating the code to simulate all the possible SynCom compositions.

## Supplementary material

**FigS1:**
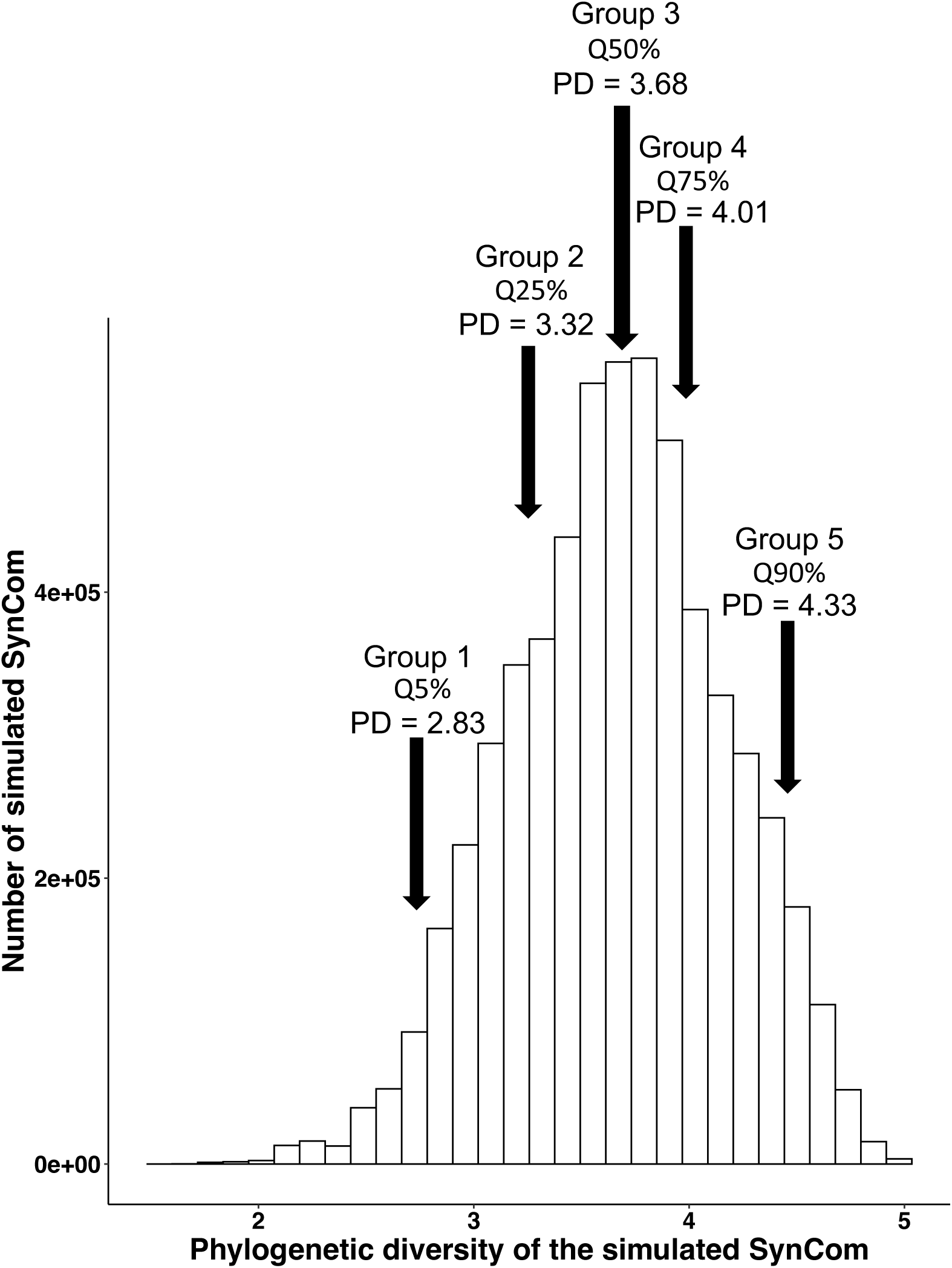
Histogram of distribution of all the simulated SynComs composed of 8 strains. The vertical black arrows indicate the quantiles selected to create the 5 groups representing a gradient of phylogenetic diversity.

**FigS2:**
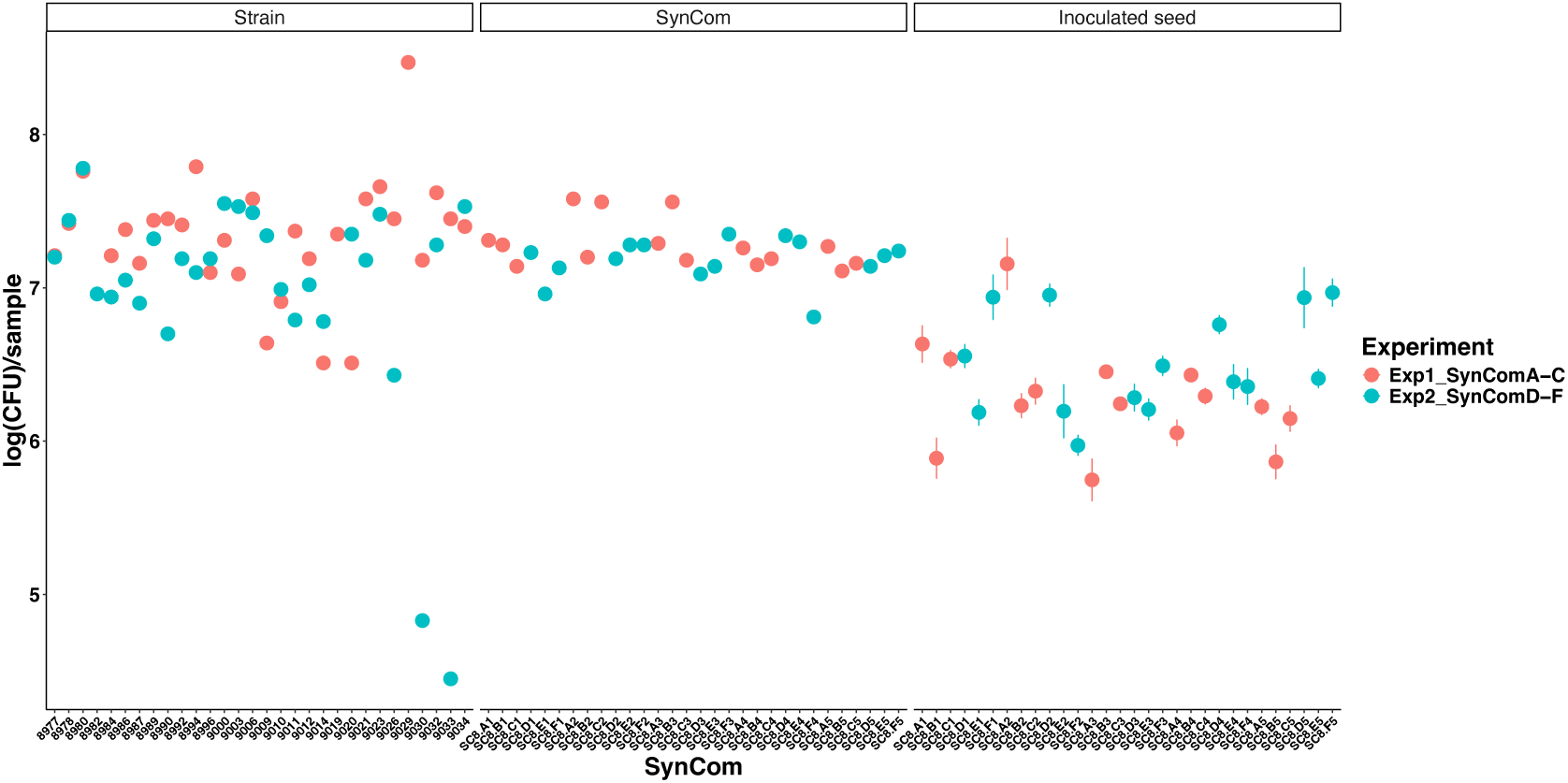
Population sizes of the inocula (individual strains and SynComs) and inoculated seeds measured as Log Colony Forming Units (CFU) per sample on TSA 10% strength plates.

**FigS3:**
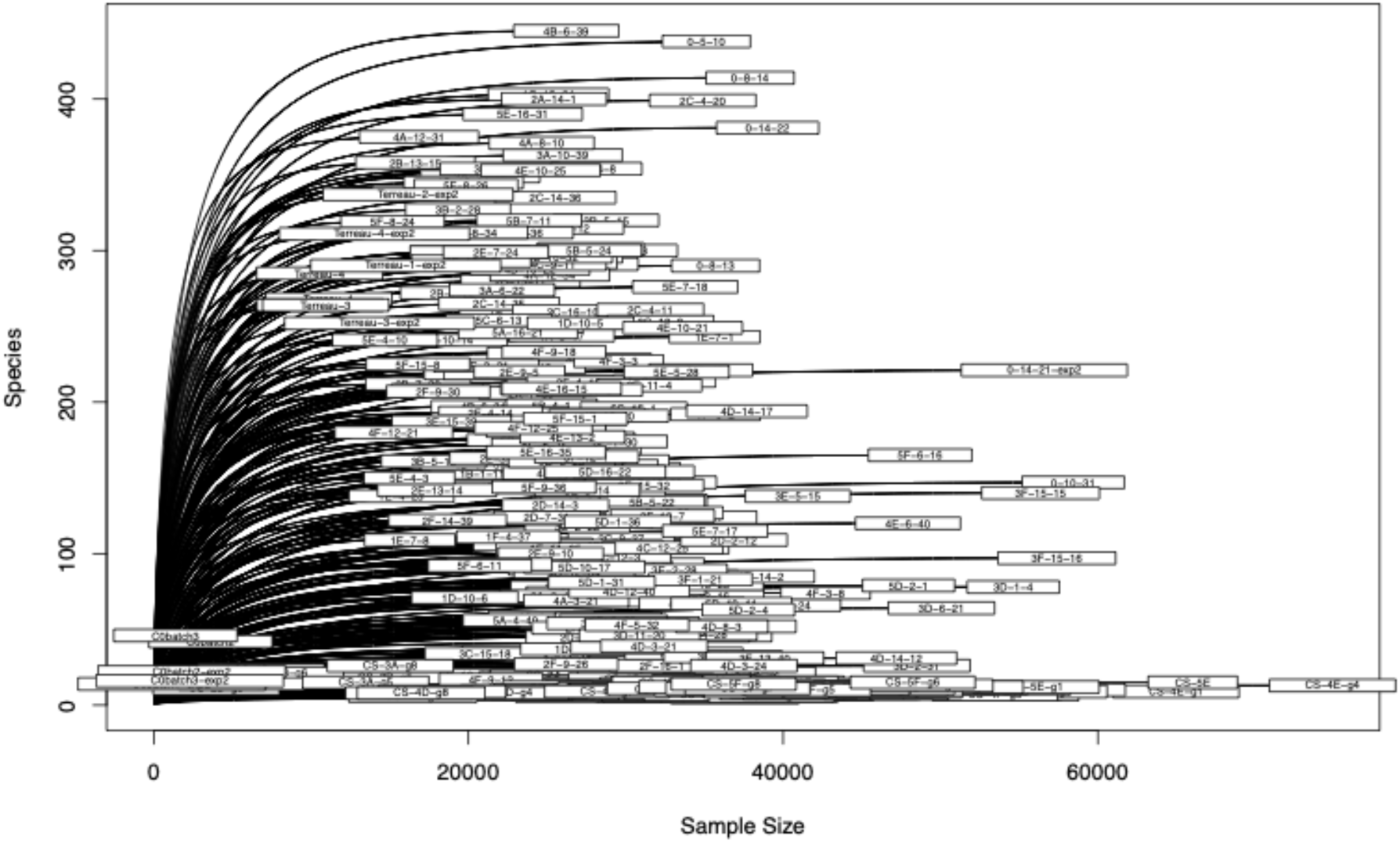
Rarefaction curves for each sample.

**FigS4:**
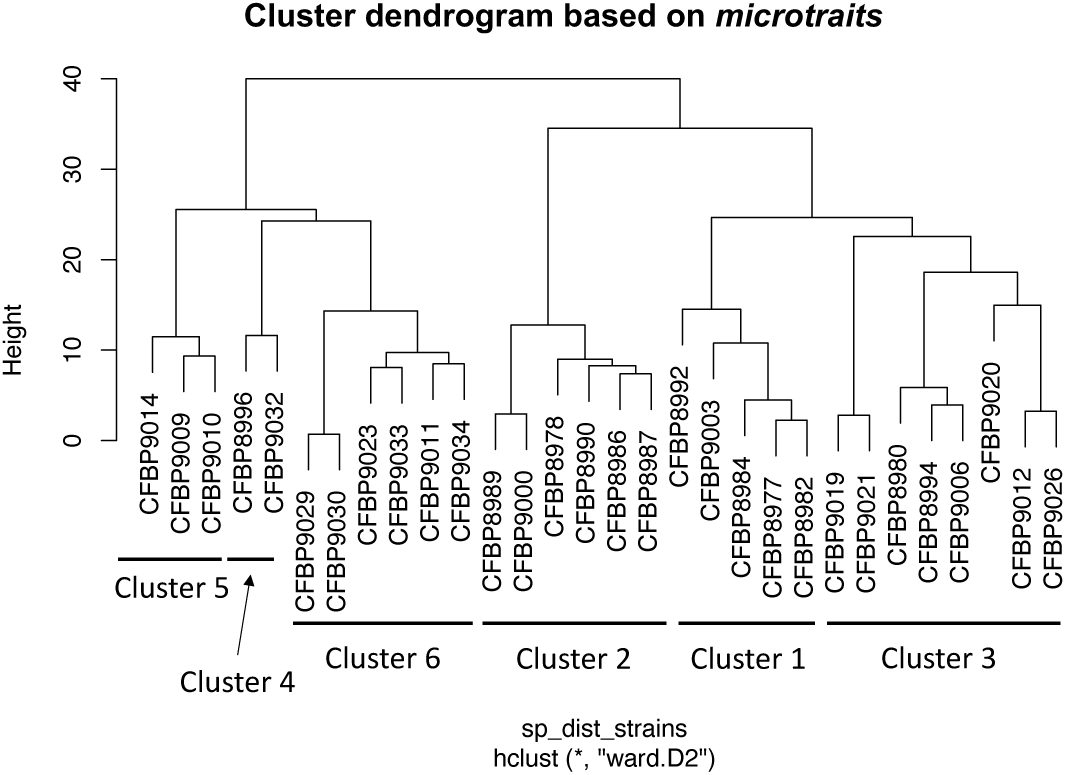
hierarchical clustering approach to group the strains in 6 clusters based on *microtraits* matrix. The matrixatgranularity3 matrix was obtained from genomes using the *microtraits* package. It consolidates strain traits into 126 distinct categories, which are further classified into four groups: resource acquisition, resource use, life history traits, and stress tolerance. From this matrix, Euclidean distances between strains were computed using the funct.dist() function from the *mFD* package. The resulting distance matrix was then used to perform hierarchical clustering (hclust()) with the ward.D2 method to obtain the 6 clusters represented.

**FigS5:**
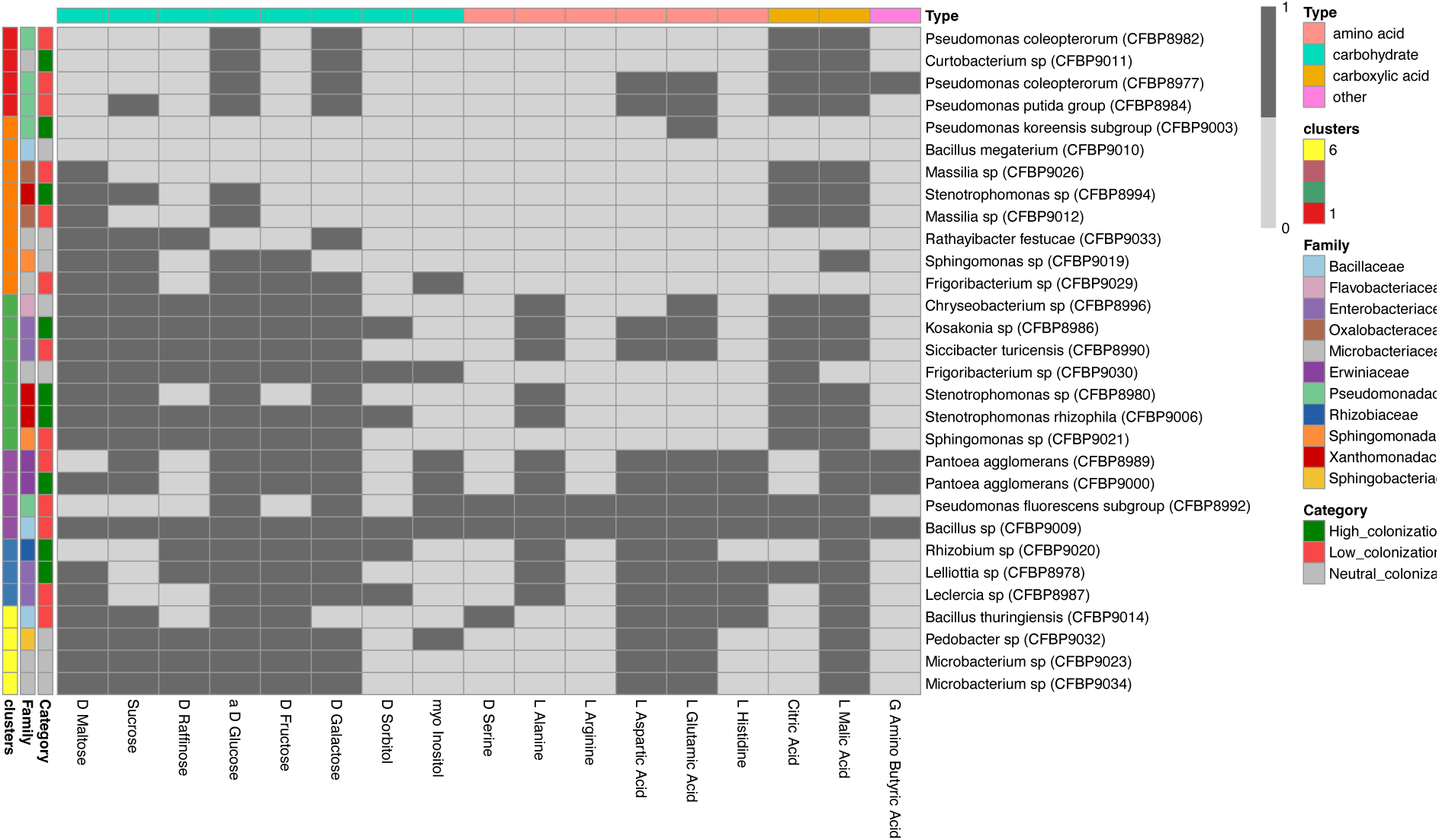
Heatmap of metabolic capacities of the strains on relevant bean seed exudates metabolites. Growing capacity of each strain on each substrate was assessed using BIOLOG plate GENIII. Dark grey box (1) indicates that strain can grow on the given substrate and light grey (0) that it cannot. The heatmap presents a subset of the 17 substrates that are found in common bean seed exudates. A hierarchical clustering approach was conducted to order the strains across 6 clusters. Substrates are colored by type (amino acid, carbohydrate, carboxylic acid and other). Strain families and colonization capacities in SynComs are indicated on the left.

**FigS6:**
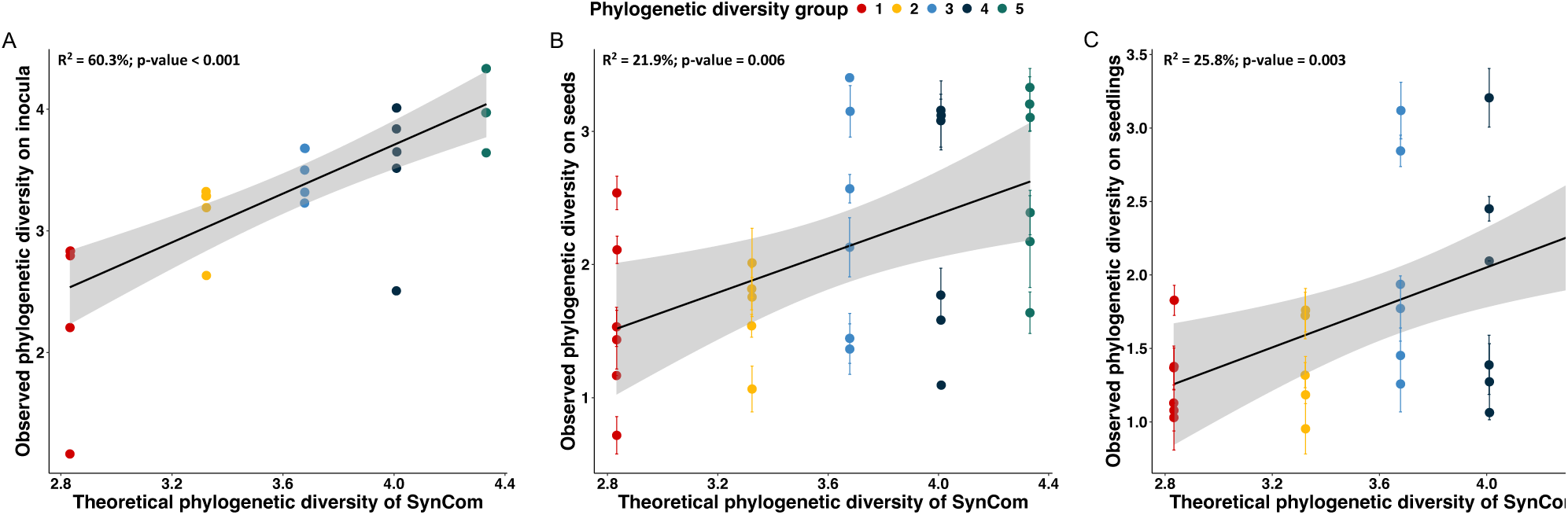
Correlations between the theoretical phylogenetic diversity (calculated as Faith’s PD) of the SynComs designed with the observed phylogenetic diversity of A) the inocula, B) the inoculated seeds, and C) the seedlings. The linear fits are drawn and the R^2^ and P-values are indicated on top of the panels.

**FigS7:**
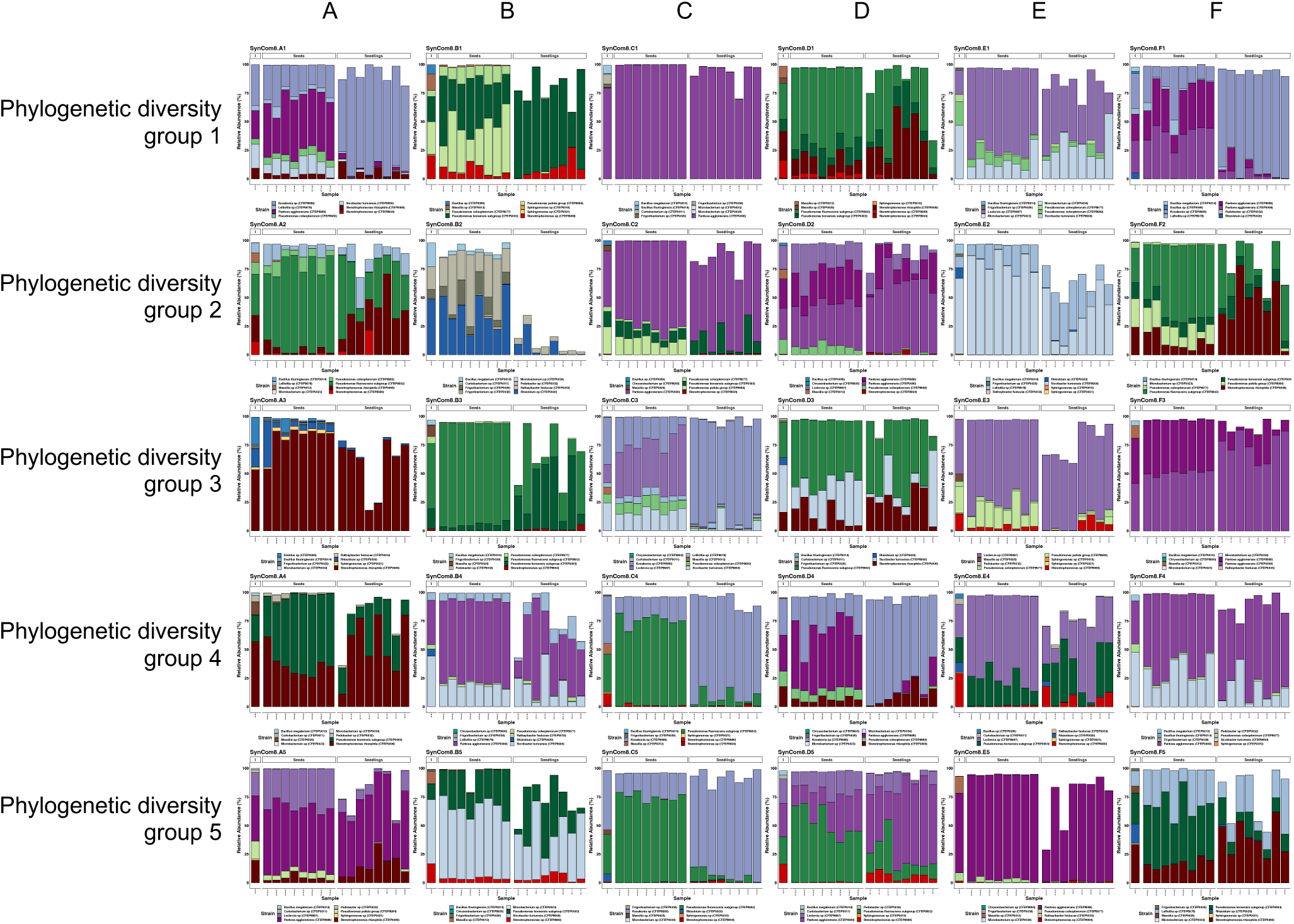
Bacterial taxonomic profiles of the inocula, inoculated seeds and seedlings for the 30 SynComs studied using *gyrB* gene metabarcoding. Only the relative abundances of the inoculated strains are displayed, the white spaces represent the contribution of other environmental bacteria (native seed, soil taxa) to the communities. Each individual bar represents a sample.

**FigS8:**
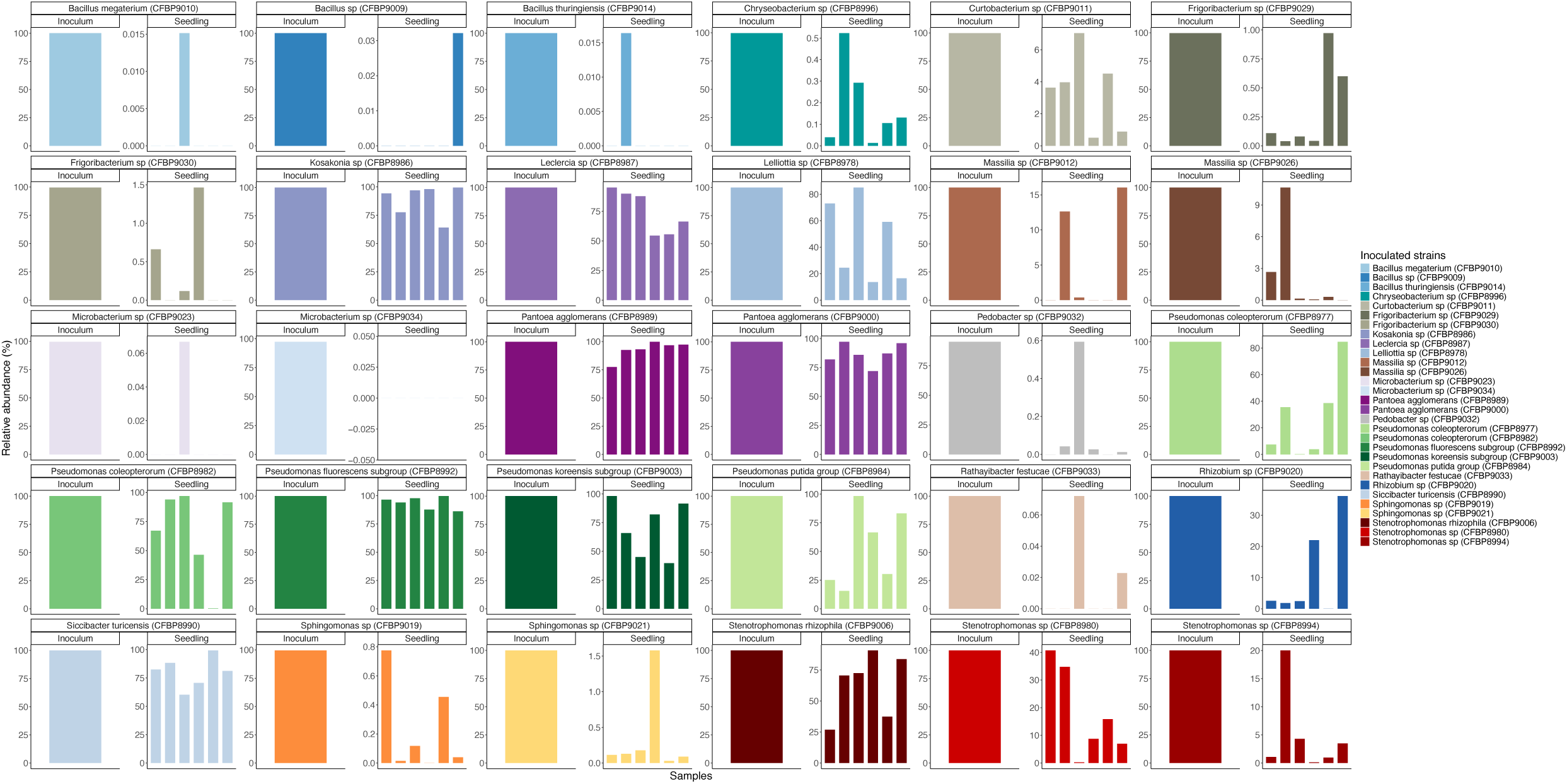
Detection of inoculated strains in both inocula and seedlings when strains are inoculated individually on seeds. The relative abundance of each strain is displayed based on gyrB gene metabarcoding. Each bar represents a sample, and the absence of a bar indicates no detection of the strain.

**FigS9:**
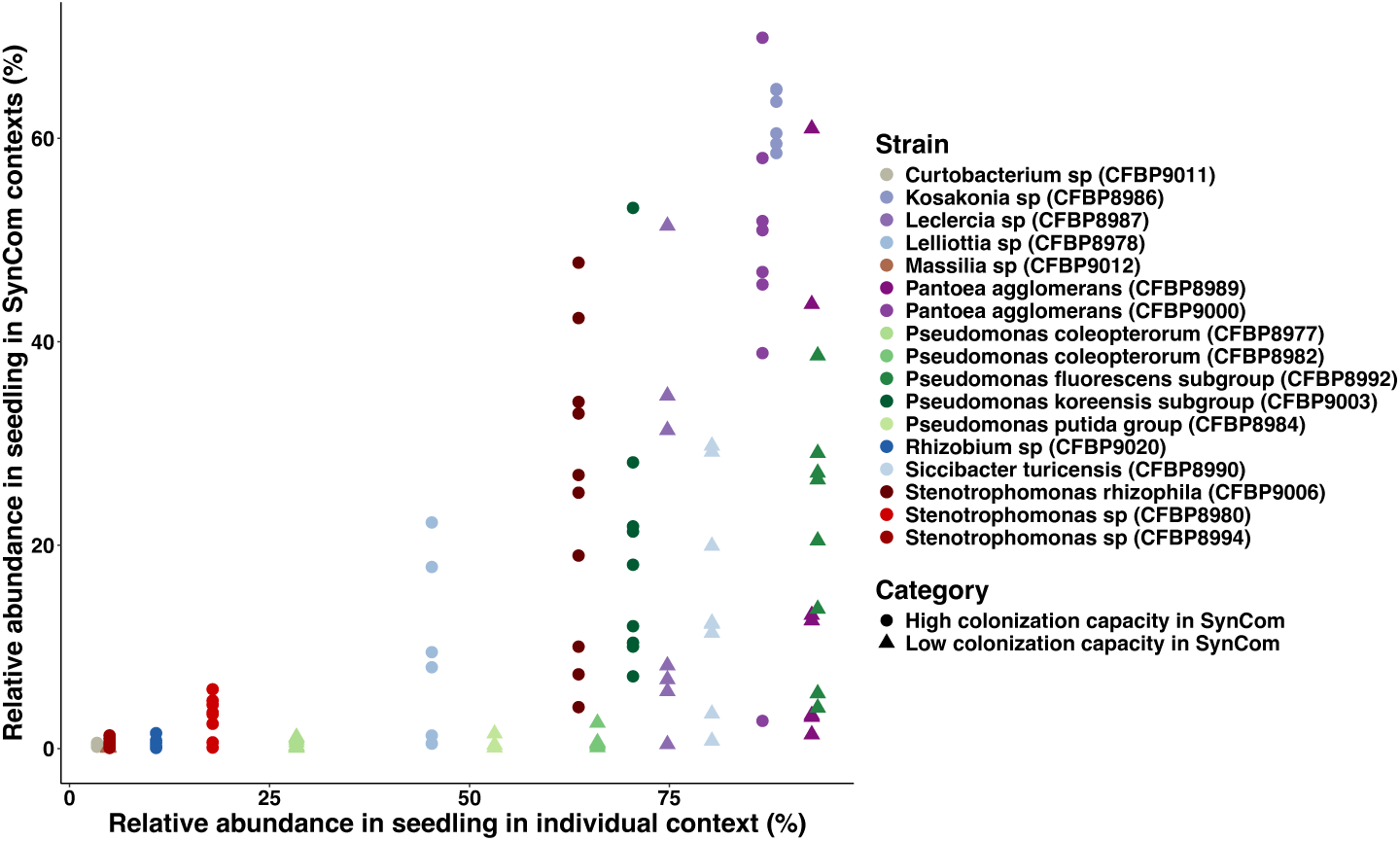
Relationship between the relative abundance of strains in seedlings in SynCom and individual contexts. For this analysis, 17 strains that were transmitted to seedling in individual inoculation and SynComs’ were kept (Mean relative abundance in seedling > 0.05%).

**FigS10:**
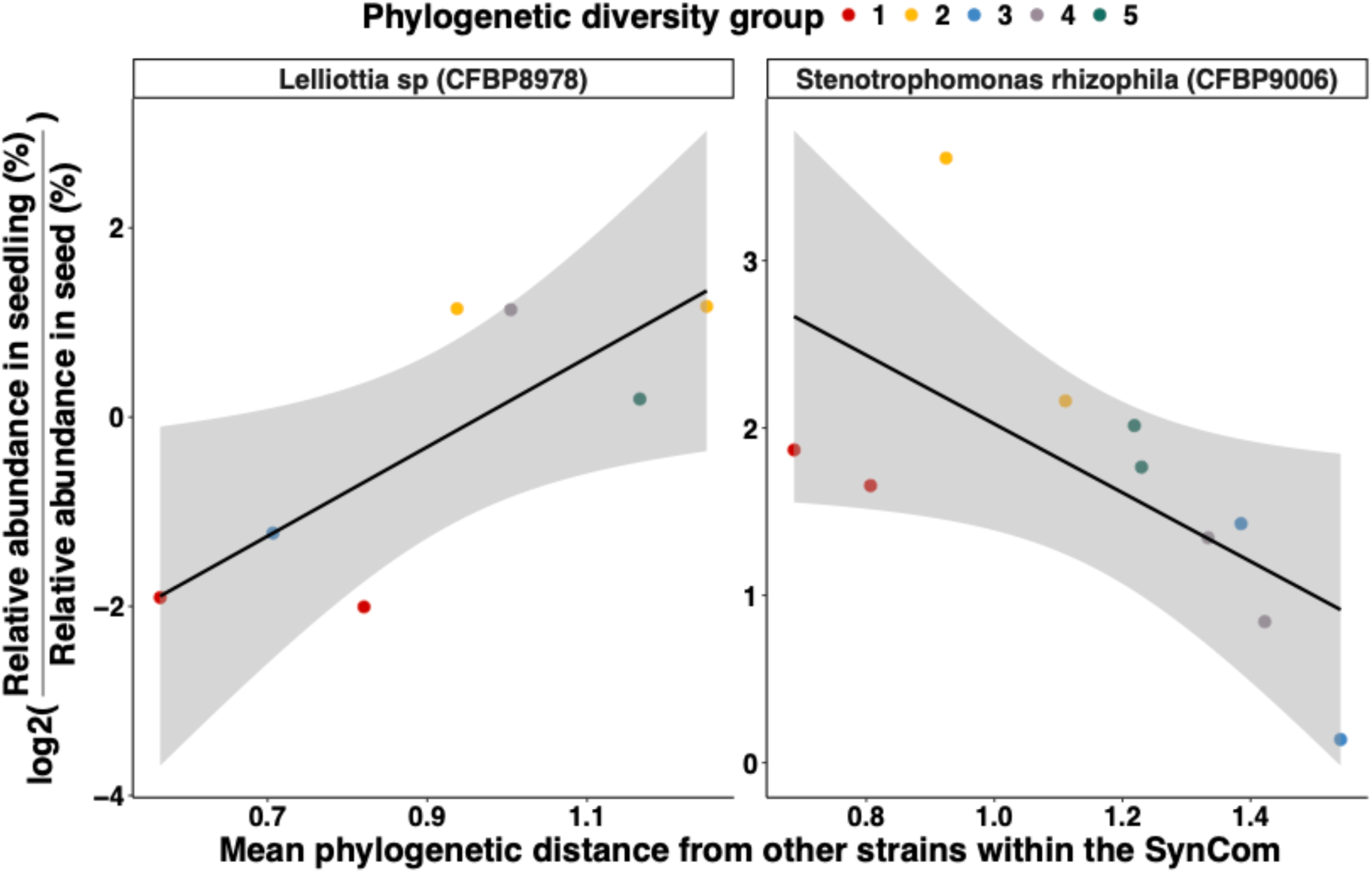
Linear regression between log2FC and mean phylogenetic distance from other strains within SynCom (MFD) for *Lellotia sp* (CFBP8978) and *Stenotrophomonas rhizophila* (CFBP9006). Linear regressions between log2FC and MFD for each strain. Only *Lellotia sp* (CFBP8978) and *Stenotrophomonas rhizophila* (CFBP9006) presented significant linear regressions (p-values < 0.05).

**Table S1:** growth capacities predicted by *gcplyr* using TSB10% growth curves.

| CFBP_strain | min_dens | max_dens | lag_time | max_percap | max_percap_time | doub_time | auc |
| --- | --- | --- | --- | --- | --- | --- | --- |
| CFBP8977 | 0,010 | 0,304 | 1,365 | 0,654 | 4,5 | 1,060 | 11,272 |
| CFBP8978 | 0,025 | 0,445 | 0,941 | 0,706 | 2 | 0,999 | 15,447 |
| CFBP8980 | 0,024 | 0,815 | 1,489 | 0,608 | 4 | 1,139 | 27,830 |
| CFBP8982 | 0,008 | 0,276 | 0,799 | 0,745 | 2 | 0,932 | 10,249 |
| CFBP8984 | 0,009 | 0,323 | 1,034 | 0,762 | 2 | 0,916 | 11,182 |
| CFBP8986 | 0,018 | 0,330 | 0,883 | 0,730 | 2 | 0,960 | 11,700 |
| CFBP8987 | 0,028 | 0,395 | 1,092 | 0,818 | 2 | 0,847 | 14,657 |
| CFBP8989 | 0,013 | 0,498 | 1,151 | 0,807 | 2,5 | 0,862 | 16,829 |
| CFBP8990 | 0,020 | 0,367 | 0,962 | 0,729 | 2 | 0,953 | 12,738 |
| CFBP8992 | 0,007 | 0,357 | 1,459 | 0,781 | 4 | 0,887 | 11,690 |
| CFBP8994 | 0,026 | 0,580 | 0,959 | 0,570 | 2 | 1,216 | 21,974 |
| CFBP8996 | 0,014 | 0,656 | 1,372 | 0,599 | 4,5 | 1,158 | 21,807 |
| CFBP9000 | 0,017 | 0,446 | 0,992 | 0,767 | 2 | 0,908 | 16,040 |
| CFBP9003 | 0,013 | 0,452 | 0,999 | 0,766 | 2 | 0,905 | 16,306 |
| CFBP9006 | 0,023 | 0,631 | 0,938 | 0,605 | 2 | 1,149 | 23,379 |
| CFBP9009 | 0,021 | 0,576 | 1,014 | 0,670 | 2,3 | 1,034 | 20,552 |
| CFBP9010 | 0,021 | 0,514 | 1,400 | 0,819 | 3 | 0,854 | 18,621 |
| CFBP9011 | 0,019 | 0,526 | 1,346 | 0,555 | 2,67 | 1,284 | 17,734 |
| CFBP9012 | 0,016 | 0,772 | 1,275 | 0,620 | 2,5 | 1,148 | 25,732 |
| CFBP9014 | 0,035 | 0,611 | 0,971 | 0,858 | 2 | 0,808 | 16,851 |
| CFBP9019 | 0,012 | 0,497 | 1,045 | 0,396 | 2 | 1,752 | 15,208 |
| CFBP9020 | 0,020 | 0,366 | 1,728 | 0,548 | 3,25 | 1,419 | 12,803 |
| CFBP9021 | 0,015 | 0,672 | 1,476 | 0,550 | 2,67 | 1,363 | 20,475 |
| CFBP9023 | 0,018 | 0,557 | 1,035 | 0,449 | 3 | 1,543 | 18,227 |
| CFBP9026 | 0,020 | 0,762 | 1,127 | 0,631 | 2,5 | 1,114 | 23,844 |
| CFBP9029 | 0,018 | 0,532 | 1,098 | 0,440 | 2,75 | 1,608 | 17,011 |
| CFBP9030 | 0,015 | 0,527 | 1,390 | 0,548 | 2,83 | 1,311 | 15,462 |
| CFBP9032 | 0,020 | 0,721 | 1,270 | 0,516 | 3,17 | 1,437 | 23,446 |
| CFBP9033 | 0,015 | 0,589 | 1,844 | 0,602 | 3,33 | 1,267 | 18,023 |
| CFBP9034 | 0,011 | 0,998 | 2,300 | 0,601 | 5,75 | 1,551 | 24,378 |

**Table S2:** Table of common bean seed exudates concentration (nmol/mL) for three replicate seed samples.

| Metabolite | Rep 1 | Rep 2 | Rep 3 | Average |
| --- | --- | --- | --- | --- |
| Saccharose | 5439 | 5851 | 8884 | 6725 |
| Citrate | 3883 | 4075 | 7107 | 5022 |
| Galactose | 1286 | 1371 | 1926 | 1528 |
| GABA | 631 | 660 | 1051 | 781 |
| Alpha-Alanine | 643 | 664 | 1029 | 778 |
| Asparagine | 608 | 592 | 1103 | 768 |
| Glutamate | 468 | 482 | 811 | 587 |
| Malate | 434 | 465 | 724 | 541 |
| Glucose | 387 | 451 | 462 | 433 |
| Arginine | 368 | 250 | 571 | 396 |
| Fructose | 317 | 326 | 363 | 335 |
| Aspartate | 231 | 237 | 412 | 293 |
| Raffinose | 251 | 253 | 344 | 283 |
| Myo-Inositol | 234 | 235 | 349 | 273 |
| Succinate | 251 | 127 | 358 | 245 |
| Xylose | 284 | 0 | 430 | 238 |
| Glycine | 155 | 170 | 271 | 199 |
| Methylcysteine | 145 | 149 | 255 | 183 |
| Threonine | 137 | 151 | 236 | 175 |
| Valine | 140 | 150 | 220 | 170 |
| Glutamine | 132 | 137 | 221 | 163 |
| Ammonium | 325 | 32 | 118 | 158 |
| Serine | 119 | 129 | 204 | 151 |
| Fumarate | 126 | 132 | 194 | 151 |
| Lysine | 120 | 123 | 186 | 143 |
| Proline | 111 | 117 | 189 | 139 |
| Leucine | 99 | 103 | 161 | 121 |
| Isoleucine | 81 | 86 | 130 | 99 |
| Tyrosine | 47 | 49 | 77 | 58 |
| Histidine | 45 | 47 | 73 | 55 |
| Phenylalanine | 45 | 46 | 68 | 53 |
| Galactinol | 33 | 30 | 51 | 38 |
| Sorbitol | 33 | 35 | 45 | 37 |
| Beta-Alanine | 31 | 32 | 47 | 37 |
| Arabinose | 30 | 35 | 45 | 37 |
| Methionine | 25 | 25 | 40 | 30 |
| Glycérate | 11 | 10 | 32 | 18 |
| Tryptophane | 14 | 16 | 22 | 18 |
| Melezitose | 7 | 7 | 11 | 8 |
| Cystine | 6 | 5 | 9 | 7 |
| Maltose | 6 | 5 | 8 | 6 |
| Ornithine | 6 | 6 | 7 | 6 |
| Melibiose | 0 | 0 | 0 | 0 |
| Trehalose | 0 | 0 | 0 | 0 |

**Table S3:** Genome informations of the different strains obtained using checkm tool (ecogenomics.github.io/CheckM/)

| CFBP | Contigs | Size | GC% | Completeness | Contamination | CDSs |
| --- | --- | --- | --- | --- | --- | --- |
| 8977 | 19 | 4820021 | 62.2 | 98.38 | 0.37 | 4265 |
| 8978 | 36 | 4442944 | 53.6 | 99.67 | 0.33 | 4033 |
| 8980 | 18 | 3975215 | 66.5 | 99.67 | 0 | 3440 |
| 8982 | 89 | 4754523 | 62.2 | 98.38 | 0.14 | 4166 |
| 8984 | 9 | 4572494 | 62.2 | 98.38 | 0.2 | 4071 |
| 8986 | 27 | 4808856 | 56.1 | 99.42 | 0.58 | 4398 |
| 8987 | 22 | 4836644 | 56.5 | 99.7 | 0.41 | 4498 |
| 8989 | 26 | 4766298 | 55.2 | 100 | 0.36 | 4375 |
| 8990 | 26 | 4131421 | 58.3 | 99.37 | 0.26 | 3759 |
| 8992 | 43 | 6313504 | 60.1 | 99.93 | 0.66 | 5710 |
| 8994 | 14 | 4250922 | 66.6 | 99.89 | 0.04 | 3717 |
| 8996 | 49 | 4439189 | 33.6 | 100 | 0 | 4011 |
| 9000 | 25 | 4725785 | 55.1 | 100 | 0.36 | 4300 |
| 9003 | 18 | 5859533 | 60.2 | 99.66 | 0.05 | 5217 |
| 9006 | 26 | 4752770 | 67.2 | 99.89 | 0.13 | 4119 |
| 9009 | 54 | 5684090 | 40.1 | 98.91 | 3.03 | 5588 |
| 9010 | 81 | 5857104 | 37.2 | 99.43 | 0.23 | 6029 |
| 9011 | 18 | 3616265 | 71.4 | 98.99 | 1.07 | 3416 |
| 9012 | 37 | 5989676 | 65.4 | 99.95 | 0 | 5147 |
| 9014 | 45 | 5836202 | 34.8 | 99.34 | 0.09 | 5769 |
| 9019 | 17 | 4245651 | 66.3 | 99.66 | 0.79 | 3791 |
| 9020 | 33 | 6136518 | 58.7 | 100 | 0.49 | 5921 |
| 9021 | 26 | 4314549 | 66.0 | 99.66 | 0.22 | 3870 |
| 9023 | 6 | 3657683 | 68.1 | 99.12 | 0.51 | 3526 |
| 9026 | 25 | 5960194 | 65.4 | 99.95 | 0 | 5122 |
| 9039 | 32 | 3208820 | 70.9 | 98.48 | 0.51 | 2937 |
| 9030 | 36 | 3266158 | 70.9 | 98.48 | 1.35 | 3008 |
| 9032 | 68 | 5582692 | 38.1 | 98.09 | 0.16 | 4687 |
| 9033 | 33 | 4341678 | 72.2 | 99.41 | 2.02 | 3915 |
| 9034 | 3 | 3691531 | 70.1 | 99.49 | 0.51 | 3346 |

**Table S4:**
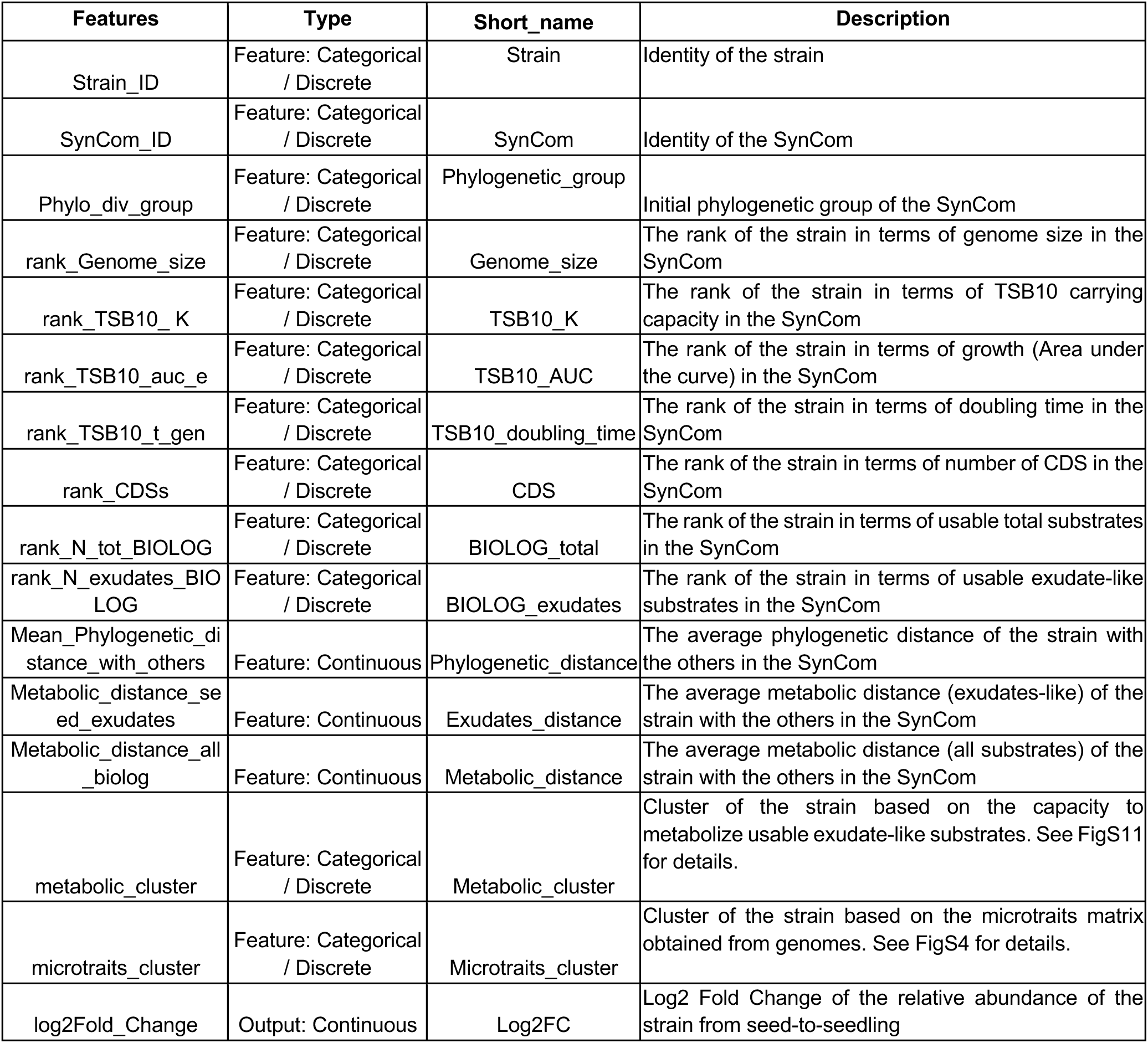
Summary of the features used for the Random Forests models.

**Table S5:**
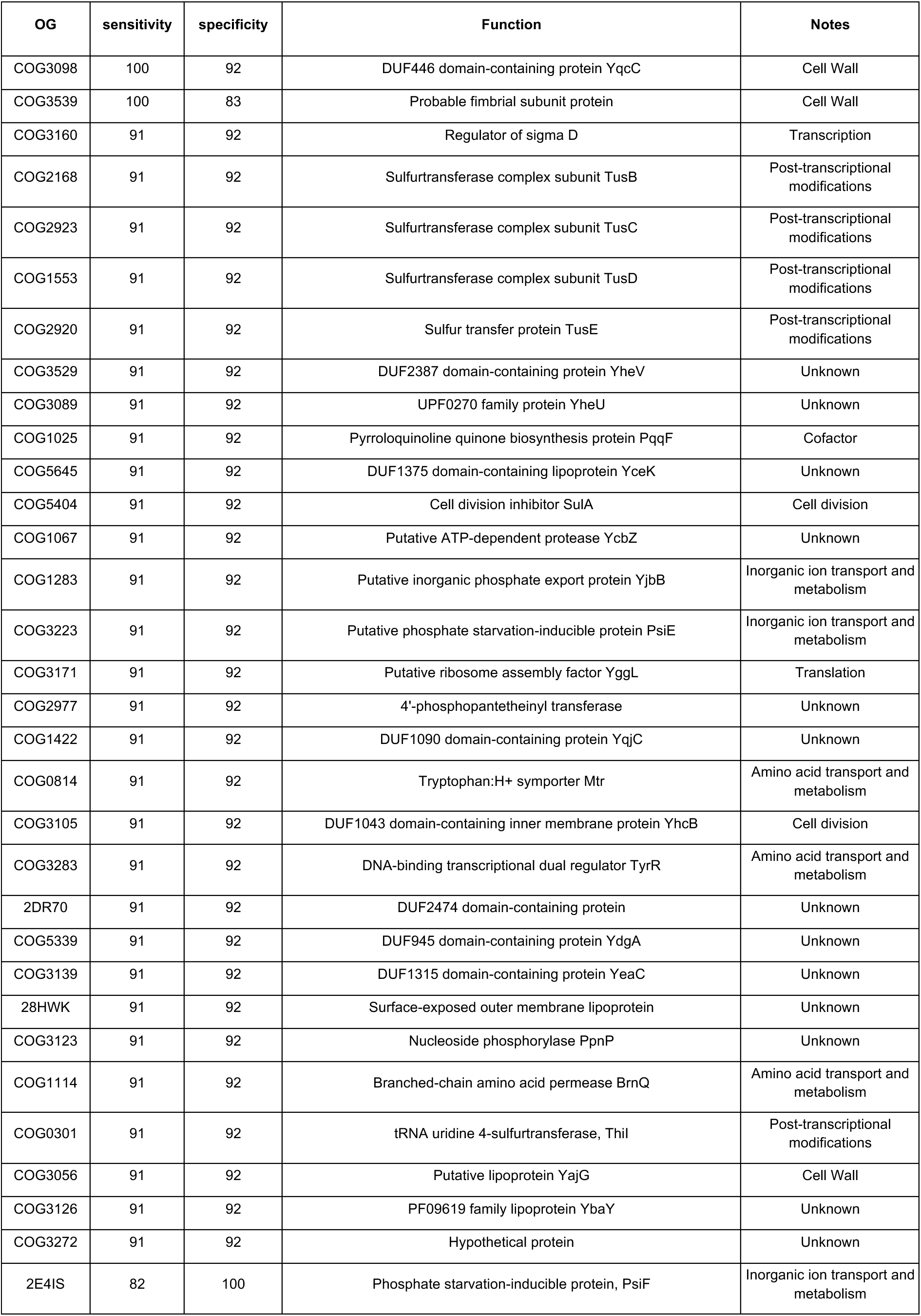
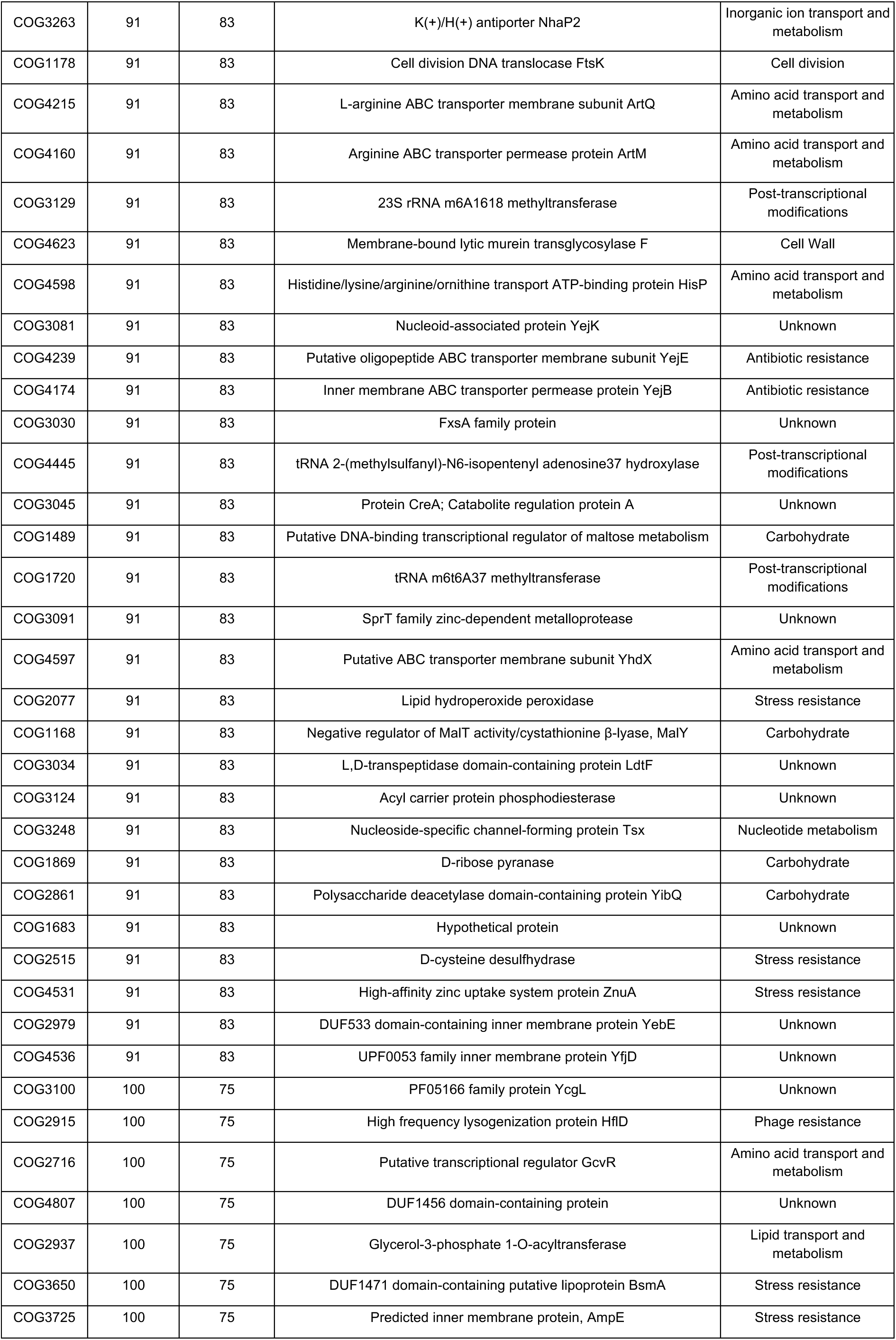
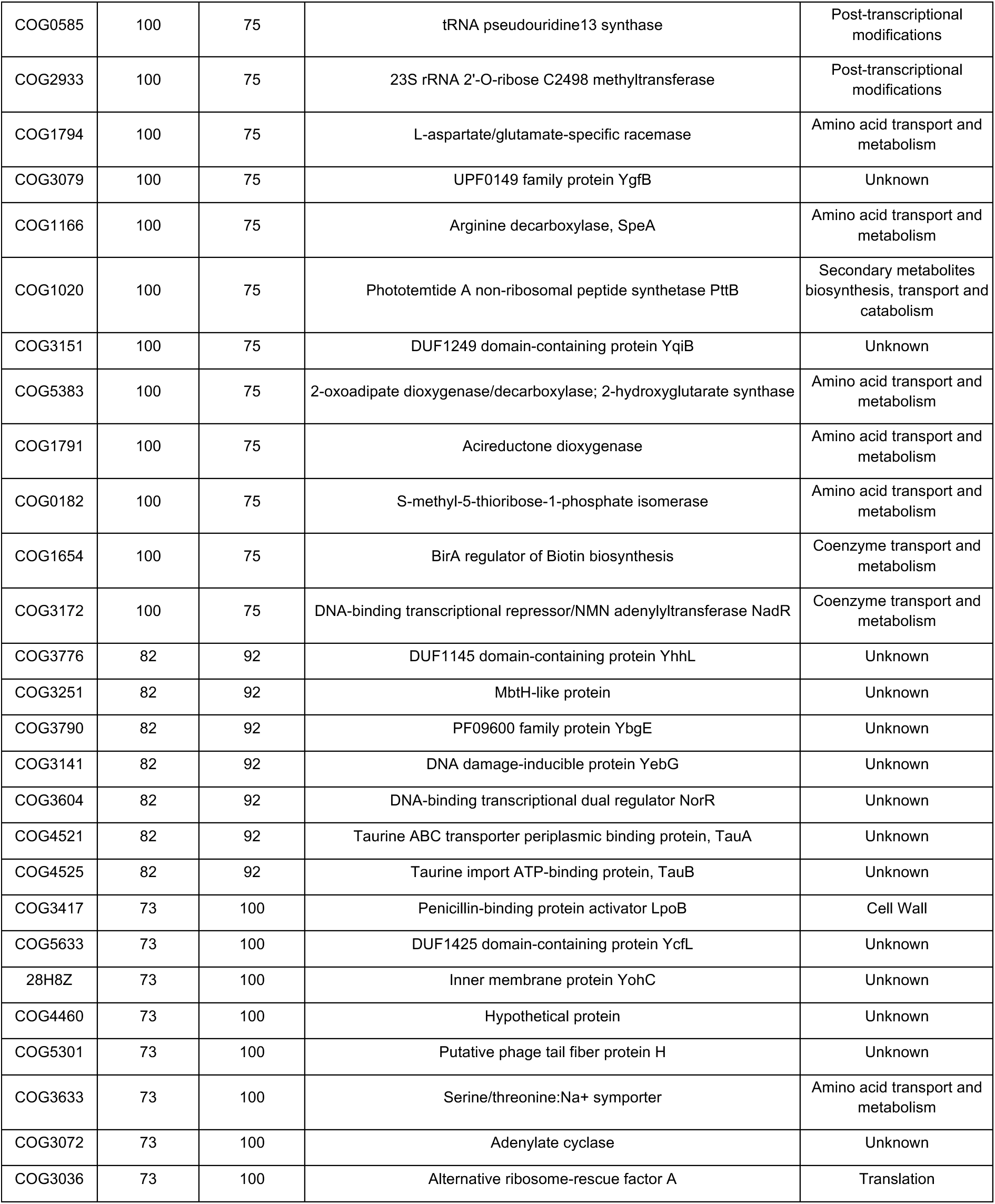
genetic traits of the high colonization success in individual contexts.

